# Biological Function Assignment Across Taxonomic Levels in Mass-Spectrometry-Based Metaproteomics via a Modified Expectation Maximization Algorithm

**DOI:** 10.1101/2025.06.12.659309

**Authors:** Gelio Alves, Aleksey Y. Ogurtsov, Yi-Kuo Yu

**Author notes:** **Corresponding Author** Yi-Kuo Yu — Division of Intramural Research, National Library of Medicine, National Institutes of Health, Bethesda, MD 20894, USA., Gelio Alves — Division of Intramural Research, National Library of Medicine, National Institutes of Health, Bethesda, MD 20894, USA.

## Abstract

A major challenge in mass-spectrometry-based metaproteomics is accurately identifying and quantifying biological functions across the full taxonomic lineage of microorganisms. This issue stems from what we refer to as the “shared confidently identified peptide problem”. To address this issue, most metaproteomics tools rely on the lowest common ancestor (LCA) algorithm to assign biological functions, which often leads to incomplete biological function assignments across the full taxonomic lineage of identified microorganisms. To overcome this limitation, we implemented an expectation-maximization (EM) algorithm, along with a biological function database, within MiCId workflow. Using synthetic datasets, our study demonstrates that the enhanced MiCId workflow achieves better control over false discoveries and improved accuracy in microorganism identification and biomass estimation compared to Unipept and MetaGOmics. Additionally, the updated MiCId offers improved accuracy and better control of false discoveries in biological function identification compared to Unipept, along with reliable computation of function abundances across the full taxonomic lineage of identified microorganisms. Reanalyzing human oral and gut microbiome datasets using the enhanced MiCId workflow, we show that the results are consistent with those reported in the original publications, which were analyzed using the Galaxy-P platform with MEGAN5 and the MetaPro-IQ approach with Unipept, respectively.

## Introduction

Mass spectrometry-based metaproteomics is a powerful high-throughput method for investigating microbiomes.^1–3^ This technique not only reveals the taxonomic composition of microbiomes but also provides insights into the relative biomass of identified taxa, reflecting the protein content of each taxon within the sample. ^4–7^ While metagenomics and metatranscriptomics shed light on the potential biological functions of microbiomes, mass spectrometry-based metaproteomics enables direct identification and quantification of these functions at a specific moment in time.^8–10^ Understanding the biological functions of microbiomes is essential, given their critical roles in human health and disease,^11,12^ agriculture,^13^ and the environment.^14,15^ For instance, the human intestinal microbiome has been implicated in conditions such as inflammatory bowel diseases, obesity, atherosclerosis, and neurodevelopmental disorders.^11^

Throughout this manuscript, we refer to biological function using the Gene Ontology (GO) framework, which categorizes biological functions into three aspects: molecular function, cellular component, and biological process. ^16^ In mass spectrometry-based metaproteomics, functional annotation begins with a set of confidently identified peptides—those identified with a proportion of false discoveries (PFD) below a defined threshold.^17,18^ These peptides are typically obtained using tandem mass spectrometry (MS/MS) database search tools such as X!Tandem,^19^ SEQUEST,^20^ and Mascot.^21^ Once identified, peptides may be used directly for functional analysis or, in some workflows, used to infer proteins, which are then utilized for functional annotation.^22^ Several bioinformatics workflows, including EggNOGmapper,^23^ MetaGOmics,^24^ Unipept,^25^ MEGAN,^26^ ProPHAnE,^27^ MetaproteomeAnalyzer,^28^ metaQuantome,^29^ and MetaX,^30^ can be used for functional and taxonomic inference in mass spectrometry-based metaproteomics.

Most workflows currently used for biological function identification in mass-spectrometry-based metaproteomics^22^ share two key characteristics: (1) confidently identified peptides are not used for microorganism identification prior to biological function identification, meaning that all protein sequences in the specified target databases are used to infer biological functions; and (2) taxonomic assignment to biological functions is based on the LCA of the confidently identified peptides or proteins. ^22^ As previous studies have shown, a consequence of not performing microorganism identification before estimating relative abundances of key quantities (such as biomass, protein, or biological function) is that confidently identified peptides may be shared by closely related taxa. Not all of these taxa may be present in the sample, or if they are, some of them are likely to be presented below the detectability range of the mass-spectrometry-based metaproteomics experiment.^31^ If not properly handled, shared confidently identified peptides can impact the accuracy of the computed relative abundances.^5,6^ Additionally, taxonomic assignment to biological functions using the LCA of confidently identified peptides does not address the shared peptide problem in mass-spectrometry-based proteomics.^32^

The shared peptide problem in mass-spectrometry-based metaproteomics is particularly prominent, as peptides are not only shared among homologous proteins within a given taxon but also between proteins from different taxa.^33–37^ We refer to this as the “shared confidently identified peptide problem” in mass-spectrometry-based metaproteomics, which arises whenever these peptides are used as shared evidence to estimate quantities of interest. This challenge can complicate the accurate assignment of biological functions and the estimation of abundances in metaproteomics analyses.

Due to the shared peptide problem encountered, the LCA of confidently identified peptides— the taxon of the most specific common ancestor—is often used to assign biological function and taxonomic composition.^24,28,38^ The LCA algorithm is a useful tool that offers a simple solution to assign biological function and taxonomic composition in metaproteomics. Additionally, the LCA algorithm is easy to implement, versatile, and, in some cases, its results can be precomputed and stored in a database for faster retrieval. However, a key limitation of the LCA approach is that it does not utilize available information from all the confidently identified peptides. For example, if 100 peptides are confidently identified, with 99 belonging to species A and 1 shared at the genus level between species A and B, the LCA algorithm would only use the evidence from the single shared peptide to estimate the quantity of interest at the genus level. It would not consider the 99 peptides from species A when making this estimate. In this case, based on the information from all the confidently identified peptides, it is apparent that species A should receive a larger fraction of the estimated quantity of interest based on the shared peptide evidence. To address this limitation, we propose an unsupervised machine learning approach—an EM algorithm—that leverages information from all confidently identified peptides to offer an alternative solution to the shared peptide problem.

In the current study, we present a modified version of the EM algorithm designed to address the challenge of using shared confidently identified peptides to compute biological function abundances across taxonomic levels. This modified algorithm builds upon the one used in our previous work for multiplexing microorganism identification from TMT-labeled samples^39^ and extends the earlier EM algorithm developed for organismal biomass estimation.^6^ While the original version was limited to biomass estimation, the modifications introduced in this generalized EM algorithm now enable the estimation of both organismal biomass and the abundances of biological functions across various taxonomic levels.

The goal of this study is to demonstrate that the newly augmented MiCId workflow— originally developed for the rapid identification of microorganisms^39–42^—can accurately identify biological functions and that the integrated, modified EM algorithm can estimate their abundances across the entire taxonomic lineage of confidently identified microorganisms in mass-spectrometry-based metaproteomics. An overview of the enhanced MiCId workflow is presented in Figure 1. To assess its performance, we evaluated the updated workflow using three synthetic datasets^29,43,44^ and two clinical microbiome datasets.^9,45^

**Figure 1.**
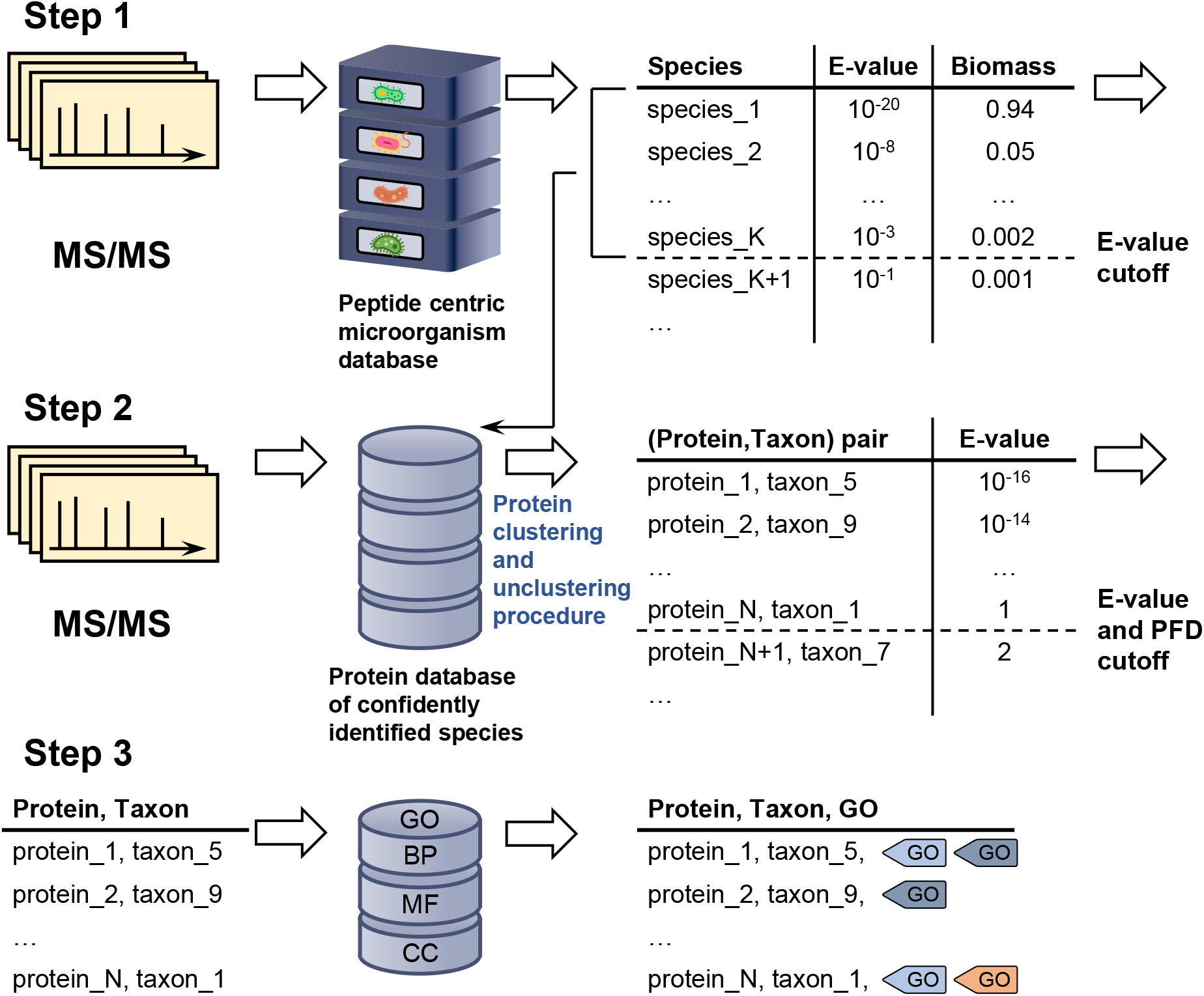
Overview of the newly augmented MiCId workflow capable to identify and compute biological function abundances in metaproteomics. The first step in this process involves querying MS/MS spectra data acquired from a sample against a comprehensive, user-defined microorganismal database built from the proteomics of microorganisms available at the NCBI database, which typically includes thousands of microorganisms. This query enables the determination of the taxonomic composition and the relative biomass of microorganisms present in the sample. In the second step, MiCId generates a protein database that includes protein sequences from reference strains of species identified with an *E*-value ≤ 0.01. The MS/MS spectra are then used to query this protein database for protein identification. During this process, a clustering procedure is applied to group proteins that share a significant number of identified peptides. Within each cluster, the best-ranking protein sequences from the identified taxa are unclustered, ensuring that only one protein sequence per taxon per cluster is retained. In the third step, protein sequences identified at a 1% proportion of false discoveries (PFD) and with an *E*-value ≤ 1 are used to query MiCId’s biological function database. At this stage, proteins, taxa, and the peptides of identified proteins are mapped to Gene Ontology (GO) terms. These GO terms, taxa, and the peptides of confidently identified proteins are used by the proposed modified expectation-maximization algorithm to compute the abundances of biological functions at different taxonomic levels. This streamlined process enables MiCId to accurately identify microorganisms, proteins, and their associated GO terms.

Using synthetic datasets, we first compared MiCId’s performance with that of Unipept and MetaGOmics in terms of microorganism identification and biomass estimation. We then assessed MiCId’s ability to identify biological functions, comparing it only to Unipept. MetaGOmics was excluded from this comparison due to its low sensitivity and high proportion of false discoveries in microorganism identification, which led to incorrect taxonomic assignments for reported biological functions. Our evaluation demonstrates that the updated MiCId outperforms both Unipept and MetaGOmics, offering improved control over the proportion of false discoveries and greater accuracy in microorganism identification and biomass estimation. In biological function identification, MiCId also shows higher accuracy and better control of the proportion of false discoveries compared to Unipept. Additionally, the modified EM algorithm in MiCId enables accurate and precise estimation of biological function abundances across the full taxonomic lineage of confidently identified microorganisms. Finally, reanalysis of clinical datasets using the enhanced MiCId workflow yielded results consistent with those reported in the original publications.

## Material and Methods

### Description of the modified EM algorithm with biomass constrain

Briefly, the presented method starts by identifying peptides in a user-specified microorganismal database. Using a set of confidently identified peptides, peptides identified with *E*-value ≤ 1, microorganismal identification is performed. Next, protein identification is carried out in a reduced database that includes the proteomes of confidently identified microorganisms at the species level. Confidently identified protein groups/clusters are then used to extract biological functions from a functional database, such as GO terms, to assign relevant biological roles to the identified proteins. In the following steps of the workflow, confidently identified peptides are mapped to their respective identified taxa and identified biological functions. For clarity, the aforementioned steps are described in Figure 1.

The proposed modified EM algorithm then calculates a probability, denoted as *p*(*t*_*α*_), for each identified taxon *t*_*α*_, using the MS^1^ extracted ion chromatogram areas of confidently identified peptides. These probabilities are constrained to sum to 1 across all identified taxa. Utilizing the computed *p*(*t*_*α*_) and the MS^1^ extracted ion chromatogram areas of confidently identified peptides, the modified EM algorithm calculates for each identified biological function *k* the quantity *p*(*k*|*t*_*α*_)*p*(*t*_*α*_), representing the joint probability of observing biological function *k* from identified taxon *t*_*α*_. A key property of these probabilities is that they are proportional to the relative biomass abundances of the taxa present in the sample.^6^ Specially, *p*(*t*_*α*_) represents the estimated relative biomass of taxon *t*_*α*_, while *p*(*k*|*t*_*α*_)*p*(*t*_*α*_) reflects the relative biomass abundance of taxon *t*_*α*_ associated with biological function *k*. Due to the biomass normalization constraint, the sum of *p*(*k*|*t*_*α*_)*p*(*t*_*α*_) over all biological functions *k* must equal *p*(*t*_*α*_), ensuring that the expected probabilities align with the relative biomassof *t*_*α*_. A detailed derivation of the computational formalism of the proposed EM algorithm, along with the protein clustering procedure, is provided in the Supporting Information file.

### MiCId’s augmented biological function database

MiCId’s workflow employs a microorganismal peptide-centric database for the rapid identification of microorganisms. Detailed information on the creation of MiCId’s microorganismal peptide-centric database has been previously described.^46,47^ As part of this study, we have enhanced MiCId’s workflow by incorporating a biological function database that links protein sequences GO terms for organisms from the NCBI database.

The augmented biological function database is constructed as follows: first, protein accession numbers from the organisms’ protein sequences (downloaded from the NCBI RefSeq database^48^) are converted to NCBI’s non-redundant protein records (WPs). Next, these WPs are used to extract GO terms from the NCBI database. The resulting GO terms, along with their corresponding WPs, form the entries in MiCId’s biological function database.

MiCId’s augmented biological function database now includes a total of 155,552,287 non-redundant protein records (WPs) with assigned GO terms. This represents 49% of the 317,321,242 WPs available in the NCBI RefSeq database, demonstrating that nearly half of these proteins have at least one GO term assigned.

### MS/MS data files (DFs)

For this study, we used three synthetic and two clinical microbiome publicly available MS/MS datasets, which were downloaded from the ProteomeXchange database at http://www.proteomexchange.^49^

The first synthetic dataset, with ProteomeXchange identifier PXD001819, contains 9 raw LC-MS/MS data files (DFs 1-9). These files represent three proteomic standard mixtures with a fixed background of yeast cell lysate spiked with 50 fmol, 25 fmol, and 12.5 fmol of the UPS1 standard protein set. Each mixture was analyzed in triplicate.^50^ The second synthetic dataset, with ProteomeXchange identifier PXD004321, includes 9 raw LC-MS/MS data files (DFs 10-18). These files correspond to three mixtures of *Staphylococcus aureus, Pseudomonas aeruginosa, Escherichia coli*, and *Streptococcus pneumoniae* combined at the following ratios: 1:1:1:1, 4:2:2:1, and 1:2:2:4. Each mixture was analyzed in triplicate.^43^ The third synthetic dataset, with ProteomeXchange identifier PXD005776, contains 3 LC-MS/MS data files (DFs 19-21) representing technical replicates of an equimolar mixture of 24 species.^44^

The first clinical microbiome dataset, with ProteomeXchange identifier PXD003151, is a human oral microbiome dataset comprising saliva and dental plaque samples from 12 children at high risk of developing dental caries. Each sample was incubated in biofilm reactors under two conditions: with sugar (WS) and without sugar (NS), as previously outlined. ^45^ In total, this dataset contains 24 samples, and for each sample two-dimensional liquid chromatography coupled with tandem mass spectrometry was conducted yielding to a total of 369 LC-MS/MS data files (DFs 22-390). This dataset was previously analyzed for biological significance^45^ and used to assess the performance of software tools in biological function analysis.^22,29^ The second clinical microbiome dataset, with ProteomeXchange identifier PXD005619, is a human gut microbiome dataset comprising stool samples from four children. Two samples are from children with ulcerative colitis (HM604, HM621), one sample from a child with Crohn’s disease in remission (HM541), and one control sample without inflammatory bowel disease (IBD) (HM609).^9^ Counting technical replicates among the samples this dataset has 9 samples, and for each sample two dimensional liquid chromatography coupled with tandem mass spectrometry was conducted yielding a total of 45 LC-MS/MS data files (DFs 391-435). Supplementary Table S1 provides pertinent information for each MS/MS file downloaded.

### MiCId and X!Tandem MS/MS data analysis

MS/MS database searches using MiCId^47^ and X!Tandem^19^ were performed with identical parameters: trypsin and Lys-C digestion allowing up to two missed cleavages. Iodoacetamide was used as a fixed modification for DFs 1–9, 19–21, and 391–435. No alkylation was applied for DFs 10–18, leaving cysteine unmodified. Methyl disulfide, also a fixed modification, was used for DFs 22–390. Mass error tolerance for precursor and product ions were extracted from the MS/MS experimental files for each dataset.

MiCId’s target databases for DFs 1–21 included protein sequences from 963,726 organisms, covering all taxa in the Unipept database. These sequences were downloaded from UniProt^51^ on February 12, 2025. For DFs 22–390 and 391–431, MiCId’s databases included proteins from 4,144 and 38,054 organisms, respectively, associated with the human oral and gut microbiomes, obtained from the NCBI BioSample database^52^ on the same date. In contrast, X!Tandem used narrower target databases limited to organisms present in the analyzed samples. Supplementary Table S2 lists all organisms included in both MiCId and X!Tandem databases.

### Unipept and MetaGOmics MS/MS data analysis

Each MS/MS data file (DFs 1-21) was initially queried using MiCId for peptide identification. Non-redundant peptides identified by MiCId, with peptide identifications controlled at a 1% PFD, were then used as input for Unipept and MetaGOmics to identify taxa and biological functions.

The Unipept search settings were configured as follows: the option to equate isoleucine and leucine was set to off, filtering of duplicate peptides was disabled, and advanced missed cleavage handling was enabled. For MetaGOmics, the search settings were configured as follows: for each DF, the protein sequences of true positive microorganisms were uploaded as a metaproteome FASTA file; for GO annotation, the UniProt sprot database was selected as the BLAST database; and for BLAST hits, a BLAST *E*-value cutoff of 1E-10 was used.

Normalized taxa abundances (NTA) were computed for each identified taxon *t*_*α*_ reported by Unipept and MetaGOmics. For each identified taxon *t*_*α*_, NTA is defined as:

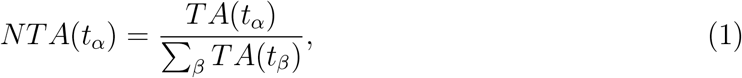

where TA(t_*α*_) represents the total number of confidently identified peptides (including redundancy) that are unique to taxon *t*_*α*_. This normalization procedure is similar to the spectrum counting normalization used for protein quantification and is also employed by some metaproteomics tools for estimating biological function abundances.^22,24,53^ Biological function abundances (BA) were computed for each GO term *g*_*α*_ reported by Unipept and MetaGOmics. The biological function abundance for a GO term *g*_*α*_ is mathematically defined as:

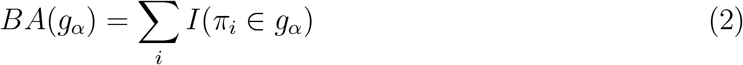

where the sum is over all confidently identified peptides (*π*_*i*_s) (including redundancy) assigned to a given GO term *g*_*α*_. Reported taxa and GO terms identified by Unipept and MetaGOmics can be found in Supplementary Tables S4-S66.

### GO term gold standard and quasi-gold standard

To properly assess the performance of the proposed EM algorithm in identifying biological functions and estimating biological function abundances at different taxonomic levels, an ideal approach would be to use a GO term gold standard dataset. However, such a gold standard is not currently available, and experimentally generating one for mass-spectrometry-based metaproteomics is a challenging task. As an alternative, we have opted to use the GO terms associated with the 48 human proteins from the UPS1 protein set as a GO term gold standard, as done in a previous study.^29^ The UPS1 protein set is a well-established benchmark designed to assess the performance of statistical methods and software in labelfree quantitative proteomics. Although it contains only 48 human proteins, these proteins map to a large number of GO terms, including 842 biological process GO terms, 125 cellular component GO terms, and 173 molecular function GO terms, making them highly suitable for our study. The full list of GO terms included in this GO term gold standard is provided in Supplementary Table S67.

Due to the lack of established GO term gold standards, we also generated what we refer to as a GO term quasi-gold standard using MS/MS data files from samples of known microbial composition, containing 4 and 24 microorganisms. For each MS/MS data file, a GO term quasi-gold standard was generated by querying the MS/MS data with X!Tandem against a target protein sequence database that contained only the protein sequences of the species present in the sample. Confidently identified proteins with *E*-values lower than the *E*-value cutoff, which controlled the expected number of false positive proteins at 1, were assigned GO terms. These GO terms were then used to create a GO term quasi-gold standard for the MS/MS data file. By restricting the target database to include only protein sequences from the species present in the sample, we ensured that the GO term quasi-gold standard contained only GO terms from true positive species. Assigning GO terms to all confidently identified proteins, without clustering, ensures broad coverage of true positive GO terms, though it may also lead to a larger number of false positive GO terms. The complete list of GO terms from the GO term quasi-gold standards is provided in Supplementary Tables S68-S94.

## Results and Discussion

### Evaluation of accuracy and precision in computed species biomass

Accurate and precise species biomass estimates in mass-spectrometry-based metaproteomics are essential for revealing microbial community composition and relative abundances. MiCId calculates biomass for confidently identified species using Eq. S10 from the proposed EM algorithm. To evaluate its accuracy, we analyzed data files from samples containing known mixtures of 4 and 24 microorganisms. The 4 microorganism samples (DFs 13–18) comprised varying proportions of the four species, while the 24 microorganism samples (DFs 19–21) contained equal amounts of each species. Fig. 2 shows box-whisker plots of the log_2_ fold change between expected biomass (EB) and computed biomass (CB) for DFs 13–21. An accurate and precise method would produce boxes centered near zero with values tightly and symmetrically distributed around it. These plots demonstrate that MiCId’s biomass estimates are both more accurate and precise than those obtained using Eq. 1 for Unipept and MetaGOmics.

**Figure 2.**
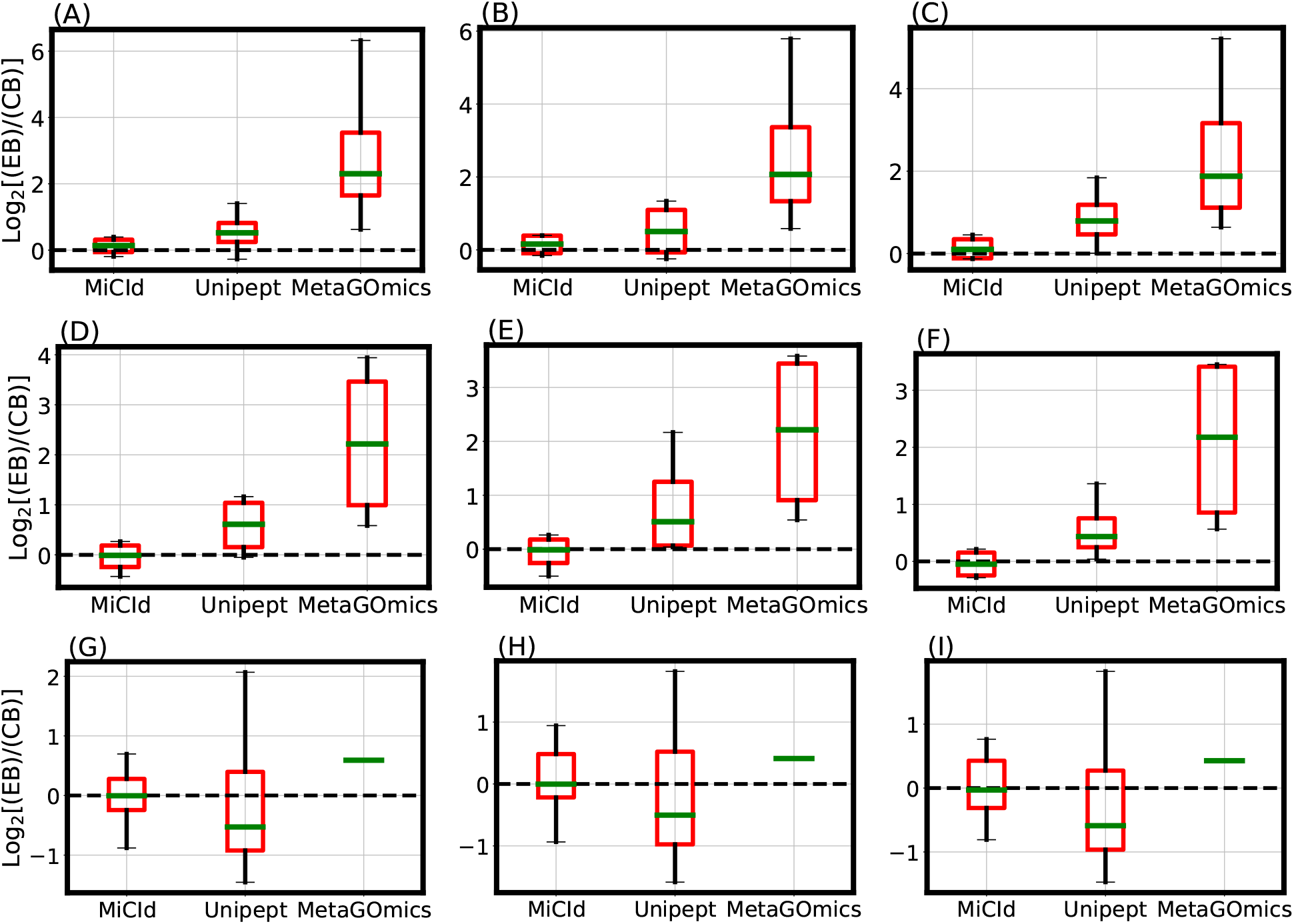
Evaluation of species level protein biomass estimates. The comparison of species level protein biomass estimates is shown through log_2_ fold change box-whisker plots representing the accuracy and precision of species biomass abundances calculated by MiCId, Unipept, and MetaGOmics. Panels (A)-(C) and (D)-(F) display the box-whisker plots from technical replicates (data files 13-15 and 16-18) for a sample composed of 4 bacterial species in a 1:2:2:4 (4:2:2:1) ratio. Panels (G)-(I) show the box-whisker plots from technical replicates (data files 19-21) for a sample composed of 24 bacterial species in a 1:1 ratio. For each sample, the log_2_ fold change statistics between the expected species biomass (EB) and the computed species biomass (CB) are displayed in the box-whisker plots. An accurate and precise biomass estimation method should produce a box-whisker plot centered around 0 with a narrow width, indicating minimal deviation from the expected biomass.

The data files used for the stacked bar plots in Fig. **??** panels (A) and (B) correspond to those in the box-whisker plots of Fig. 2 panels (A) and (G), respectively. While the box-whisker plots display computed biomass for true positive species only, the stacked bar plots include both true and false positives. Panel (A) shows results for DF 13, a mixture of *Staphylococcus aureus, Pseudomonas aeruginosa, Escherichia coli*, and *Streptococcus pneumoniae* in a 4:2:2:1 ratio. MiCId’s biomass estimates closely match the expected values, correctly identifying all four true positive species without false positives. In contrast, Unipept and MetaGOmics show greater deviation: Unipept identified all four true positives but also 15 false positives, while MetaGOmics identified all four true positives along with 27 false positives. Panel (B) shows results for DF 19, a 24-species equal-ratio mixture. Here, MiCId’s biomass estimates align better with expectations, and it correctly identifies 21 of 24 true positives with only 3 false positives. Unipept identified 16 true positives with 2 false positives, and MetaGOmics identified just 1 true positive with 21 false positives. Similar trends were observed for data files 10–12, 14–18, and 20–21 (Supplementary Figs. S1–S4).

Accurate and precise biomass estimation by the proposed EM algorithm depends on high microorganism identification sensitivity and effective false positive control. Table 1 summarizes true and false positive species identified by MiCId, Unipept, and MetaGOmics, both without and with false positive control where applicable. MiCId controls false positives using an *E*-value cutoff of 0.01, excluding identifications above this threshold. For Unipept, we applied a filter removing identifications representing less than 0.5% of taxon-specific peptides.^54^ With false positive control applied, MiCId identified 97 true positives and 10 false positives out of 108 possible species, achieving 90% sensitivity and a 9.3% PFD. Unipept identified 84 true positives and 102 false positives (77.7% sensitivity, 54.8% PFD), while MetaGOmics identified 36 true positives and 325 false positives (33.3% sensitivity, 90% PFD).

**Table 1:** Performance evaluation of species level identification of MiCId, Unipept, and MetaGOmics. Samples of mixtures composed of 4 microorganisms at varying ratios (DF 10-18) and mixtures composed of 24 microorganisms at equal ratios (DF 19-21) were used for the performance evaluation. All peptides identified by MiCId, with the proportion of false discoveries (PFD) controlled at 1%, were used as input for MiCId, Unipept and MetaGOmics. The data files (DF) used are listed in the first column. For the MiCId and Unipept results, subscripts “u” (or “f”) for true positives (TP) and false positives (FP) indicate whether the filtering strategy to control false positives was turned off (or on). In MiCId results, “u” includes all identified taxa with an *E*-value ≤ 1, while “f” includes only taxa with an *E*-value ≤ 0.01. For Unipept, the filtering strategy described in ^54^ was applied, where any species identified with less than 0.5% of the total number of taxon-specific peptides identified were considered true negatives and removed from the analysis. For MetaGOmics, no filtering strategy was applied, as we are unaware of any recommended approach to control the number of false positives specific to MetaGOmics.

| Species level identifications for data files 10-21 |  |  |  |  |  |  |  |  |  |  |
| --- | --- | --- | --- | --- | --- | --- | --- | --- | --- | --- |
| Data File | MiCId |  |  |  | Unipept |  |  |  | MetaGOmics |  |
|  | TP<sub>f</sub> | FP<sub>f</sub> | TP<sub>u</sub> | FP<sub>u</sub> | TP<sub>f</sub> | FP<sub>f</sub> | TP<sub>u</sub> | FP<sub>u</sub> | TP<sub>u</sub> | FP<sub>u</sub> |
| 10 | 4 | 0 | 4 | 0 | 4 | 5 | 4 | 102 | 3 | 25 |
| 11 | 4 | 0 | 4 | 0 | 4 | 6 | 4 | 121 | 3 | 25 |
| 12 | 4 | 0 | 4 | 1 | 4 | 8 | 4 | 109 | 3 | 26 |
| 13 | 4 | 0 | 4 | 0 | 4 | 15 | 4 | 306 | 4 | 27 |
| 14 | 4 | 0 | 4 | 0 | 4 | 10 | 4 | 320 | 4 | 29 |
| 15 | 4 | 0 | 4 | 0 | 4 | 15 | 4 | 345 | 4 | 29 |
| 16 | 4 | 0 | 4 | 0 | 4 | 8 | 4 | 265 | 4 | 30 |
| 17 | 4 | 0 | 4 | 0 | 4 | 13 | 4 | 324 | 4 | 32 |
| 18 | 4 | 0 | 4 | 0 | 4 | 17 | 4 | 180 | 4 | 32 |
| 19 | 21 | 3 | 21 | 3 | 16 | 2 | 24 | 573 | 1 | 21 |
| 20 | 20 | 4 | 20 | 4 | 16 | 1 | 23 | 589 | 1 | 24 |
| 21 | 20 | 3 | 20 | 3 | 16 | 2 | 22 | 632 | 1 | 25 |

Notably, among the false positives identified by MiCId and Unipept for DFs 19–21 (which include 24 true positive species), were *Salmonella enterica, Escherichia coli*, and *Bacillus spizizenii*. These were misidentified in place of, or co-identified with, the true positives *Salmonella bongori, Shigella flexneri*, and *Bacillus subtilis*, respectively. Such misidentifications highlight a common challenge in metaproteomics when using large microorganismal databases—here comprising 963,726 organisms—where closely related taxa can be incorrectly or jointly identified. Additionally, Table 1 shows that Unipept’s heuristic filtering strategy ^54^ does not consistently limit false positives, whereas MiCId’s *E*-value thresholding is more effective. No filtering strategy was applied to MetaGOmics, as no standard method for false positive control is available. The full list of false positives is provided in Supplementary Table S3.

Accurate biomass estimation of identified microorganisms is critical for MiCId, as these values serve as constraints in the EM algorithm for computing biological function abundances across the full taxonomic lineage. MiCId’s species level sensitivity and PFD are consistent with prior studies using larger, more complex datasets.^6,47^ The results also highlight the need for improved statistical methods to control the PFD in taxonomic assignments when using LCA-based approaches, in line with earlier findings.^55^ Furthermore, refining biomass estimation—particularly when derived from confidently identified, taxon-specific peptides—remains important, especially in addressing redundancy. MetaGOmics was excluded from further analysis due to its low sensitivity and PFD, which hindered accurate functional assignment to true-positive species, the primary goal of this manuscript. Complete MetaGOmics results for the synthetic datasets are available in Supplementary Tables S25–S66.

### Assessment of GO term identification sensitivity and proportion of false discoveries (PFD)

Panel (A) of Fig. 3 shows a Venn diagram comparing biological process GO terms identified by MiCId, Unipept, and X!Tandem with the UPS1 GO term gold standard, illustrating both unique and shared GO terms. An accompanying overlap coefficient matrix quantifies the fraction of shared GO terms relative to each method’s total. When including the gold standard, the matrix row indicates GO term identification sensitivity, while one minus the values in the gold standard column represents the PFD (i.e., 1-precision). From the matrix, X!Tandem achieves 100% sensitivity (842/842) but a high PFD of 38.3% (522/1364). MiCId shows 94.8% sensitivity (798/842) with a low PFD of 8.6% (75/873). Unipept has lower sensitivity (24%, 198/842) and a high PFD (81.8%, 892/1090). Similar trends hold for molecular function and cellular component GO terms, with detailed Venn diagrams and matrices available in Supplementary Fig. S5 and replicates DFs 2–3 shown in Supplementary Figs. S6 and S7.

**Figure 3.**
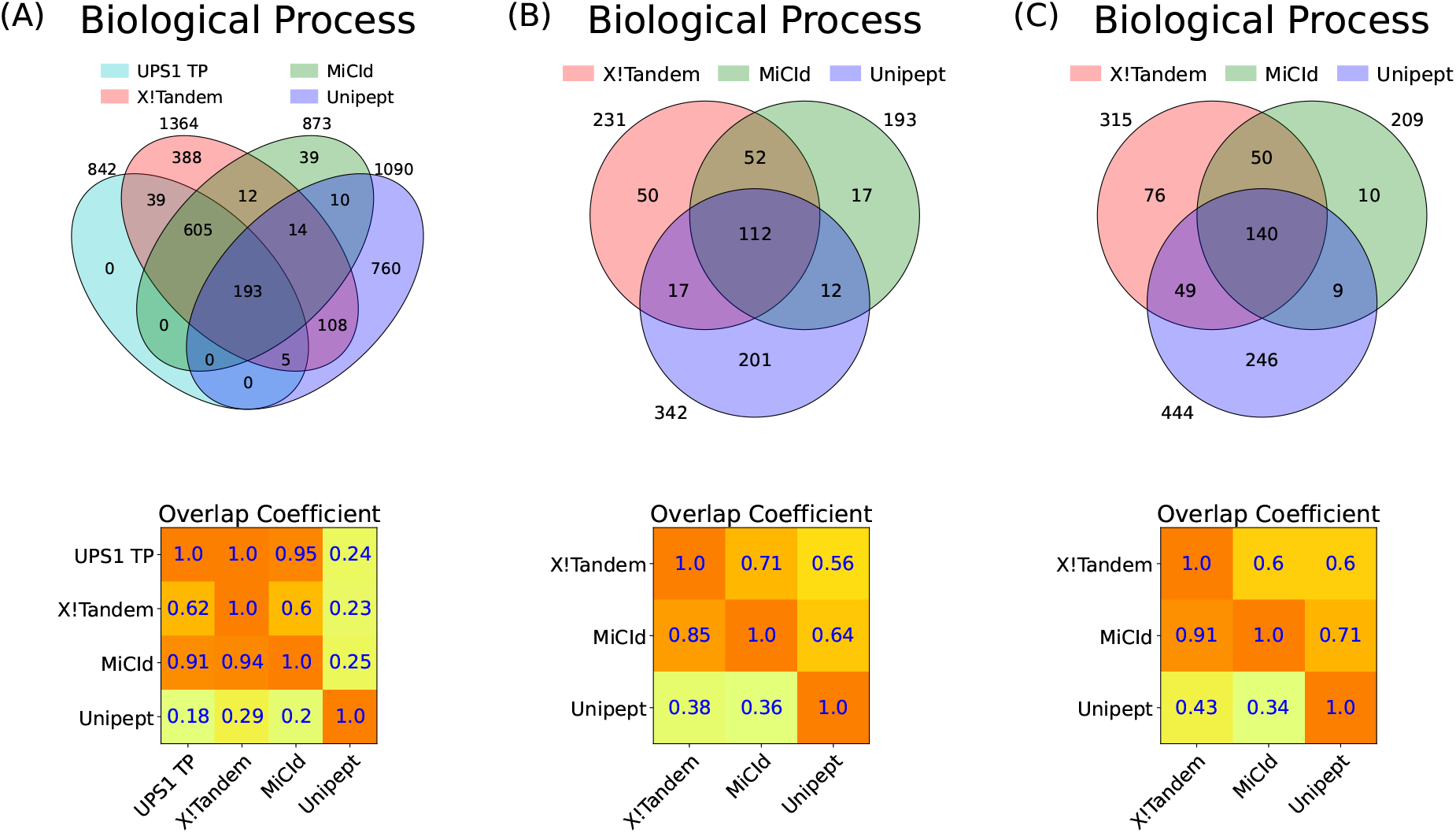
Assessment of GO term identification sensitivity and the proportion of false discoveries (PFD) is demonstrated through the analysis of mixtures from 48 human proteins from the UPS1 dataset in panel (A), 4 microorganisms in panel (B), and 24 microorganisms in panel (C). Panels (A)-(C) present Venn diagrams and an overlap coefficient matrix for the GO terms reported by X!Tandem, MiCId, and Unipept. Panel (A) also includes the GO terms from the gold standard for the 48 human proteins in the UPS1 protein set (UPS1 TP). The Venn diagrams illustrate the number of GO terms identified by each method, as well as the number of terms co-identified by the methods. The overlap coefficient matrix values are calculated as the fraction of GO terms in the intersection between methods (corresponding to the row and column), divided by the total number of GO terms reported by the method in the row. When a GO term gold standard is used, as in panel (A) for UPS1 TP, the values in the first row of the matrix represent the sensitivity of the methods listed, while one minus the values in the first column indicates the PFD for those methods. In panels (B) and (C), where a GO term quasi-gold standard is used (based on the GO terms reported by X!Tandem), the magnitude of the values in the first column reflects the performance of the methods listed in that column. The GO terms identified from the analysis of data files 7, 10, and 19 are used in panels (A), (B), and (C), respectively.

To evaluate GO term identification in multi-microorganism samples, we used GO term quasi-gold standards for data files 10–12 and 19–21, derived from MS/MS analyses of mixtures containing 4 and 24 microorganisms, respectively. Unlike a true gold standard, the overlap coefficient matrix’s row and column containing a quasi-gold standard do not represent sensitivity and PFD in the same way. Instead, values in the quasi-gold standard column indicate how well each method identifies true positive GO terms, as the quasi-gold standard includes all GO terms linked to confidently identified proteins without clustering. This set likely contains many true positives, along with some false positives. Panel (A) of Fig. 3 supports this, X!Tandem identified all true positives but also reported 522 false positives. Approximately 94% of MiCId’s identified GO terms overlap with X!Tandem’s, reflecting high specificity and only 9% false positives, indicating that MiCId’s large overlap with X!Tandem reflects mostly true positives. In contrast, Unipept’s low sensitivity and high false positive rate mean its overlap with X!Tandem likely stems from false positives. Supplementary Figs. S5–S7 corroborate these findings. Panels (B) and (C) of Fig. 3 show that MiCId outperforms Unipept in multi-species samples, with an average overlap coefficient of 88% versus 44% with X!Tandem. Similar trends are seen for molecular function and cellular component GO terms (Supplementary Figs. S8, S11) and for replicates of DFs 10 and 19 (Supplementary Figs. S9, S10, S12, S13).

These results demonstrate that the newly implemented protein clustering procedure in MiCId, detailed in the Supporting Information File, is functioning as intended. The low PFD indicates that MiCId’s clustering procedure effectively groups proteins, minimizing false positives. In contrast, the high PFD observed with X!Tandem highlights the limitations of not using a clustering procedure for identified proteins. Moreover, the high sensitivity for reported GO terms in MiCId shows that its unclustering procedure is successfully separating the correct proteins of confidently identified species. It is also important to note that having accurate statistical significance in terms of *E*-values^40^ assigned to identified proteins in MiCId’s workflow plays a crucial role in both the clustering and unclustering procedures. The *E*-values provide a robust measure to rank identified proteins within clusters and to effectively control the PFD of identified proteins,^17,40^ all without requiring a decoy database. Ultimately, it is through the confidently identified protein heads of clusters that GO terms are retrieved from MiCId’s ontology database and assigned to identified species.

### Quantitative analysis of computed GO term abundances

Accurately computing GO term abundances at each taxonomic level is essential for understanding how microbial community functions shift in response to perturbations. To assess the accuracy of computed GO term abundances, we analyzed MS/MS data from DFs 1–3, 4–6, and 7–9, which contain 48 human UPS1 proteins spiked into yeast lysate at concentrations of 12.5, 25, and 50 pmol, respectively. Panels (A), (B), and (C) in Fig. 4 show box-whisker plots of the log_2_ fold changes for protein ratios of 1:1, 2:1, and 4:1. Panel (A) shows that MiCId’s log_2_ fold change values are tightly centered around the expected value of zero, indicating both high accuracy and precision in computing biological process GO term abundances (Eq. S13). Unipept’s values are similarly centered but more widely distributed, suggesting comparable accuracy but lower precision (Eq. 2). Panels (B) and (C) show Mi-CId’s average log_2_ fold changes close to the expected values of 1 and 2, with slight skewing indicating minor precision loss. In contrast, Unipept’s values deviate from expectations, reflecting lower accuracy. Similar trends were observed for cellular component and molecular function GO term abundances (Supplementary Figs. S14 and S15).

**Figure 4.**
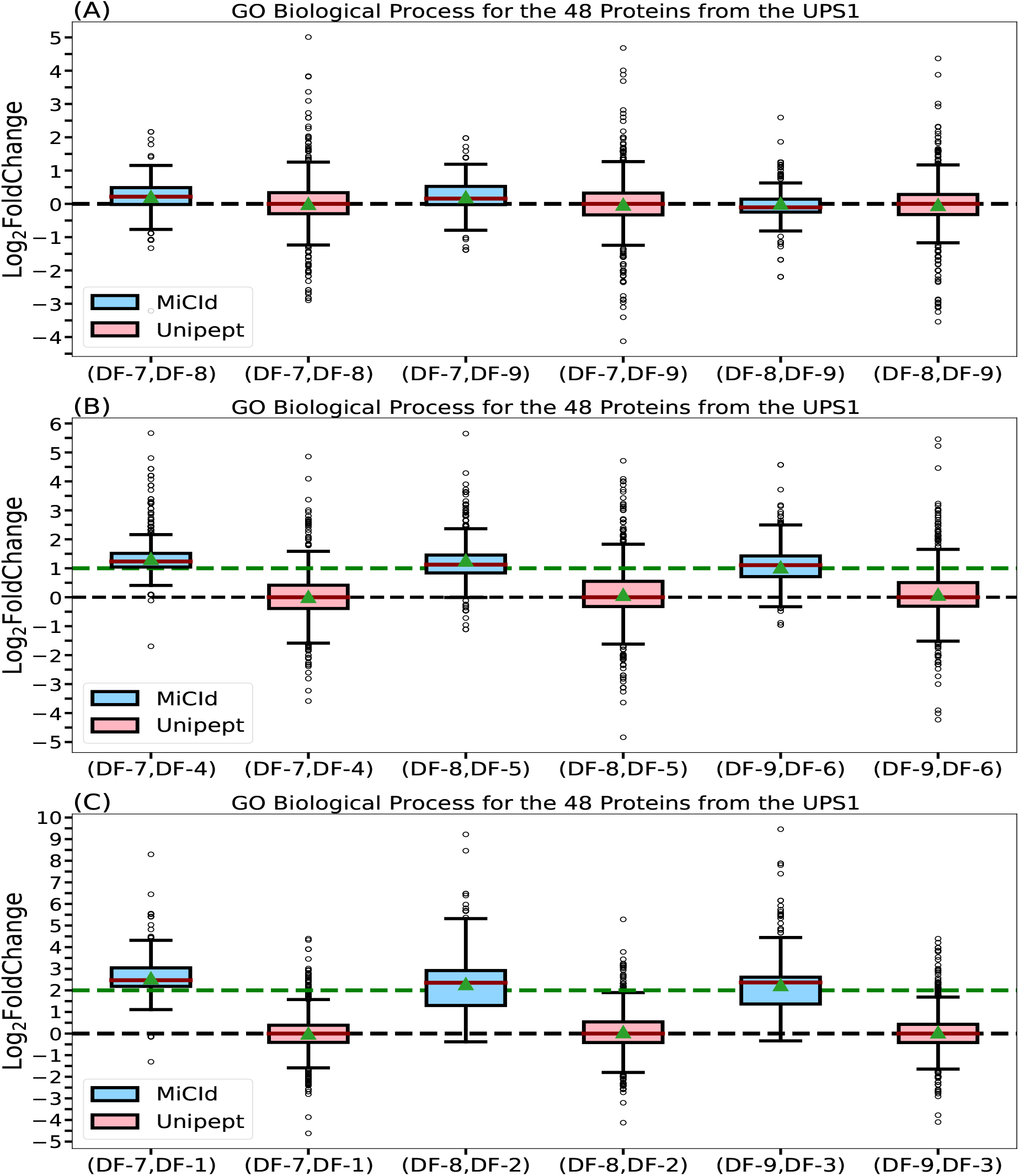
Assessment of GO term abundances computed by MiCId and Unipept. Panels (A), (B), and (C) present box-whisker plots for the log_2_ fold change between samples with varying amounts of the 48 human proteins from the UPS1 protein set, at 1:1, 2:1, and 4:1 ratios, respectively. In each panel, an accurate and precise biomass estimation method should result in a box-whisker plot centered around the expected values of 0, 1, and 2, respectively, with a narrow width, indicating minimal deviation from the expected biomass. The values for median and average biomass are represented by a solid red line and a green triangle, respectively, in the box-whisker plots. The log_2_ fold change in GO term abundances is derived from the analysis using the data files (DF-i, DF-j), as indicated by the x-axis labels for each box-whisker plot.

The outliers observed in panels (B) and (C) of the log_2_ fold change box-whisker plots for both MiCId and Unipept primarily result from missing data.^56,57^ These missing values arise due to differences in protein identification confidence across samples with varying UPS1 concentrations. Higher protein amounts lead to more confident identifications and thus higher GO term abundances. In our dataset, samples with 50, 25, and 12.5 pmol of UPS1 typically yielded identifications of 48, 40, and 34 proteins, respectively, with varying peptide counts. Consequently, when computing log_2_ fold changes between samples with different UPS1 concentrations, outliers skewed toward higher values are expected, as the GO term abundances are derived from different protein sets. These outliers do not reflect inaccuracies in the EM algorithm’s computed abundances but rather highlight the impact of missing data—a challenge the EM algorithm was not designed to address. As shown in panel (A) of Fig. 4 and Supplementary Figs. S14 and S15, this effect is minimal when comparing technical replicates. Therefore, subsequent evaluations of the EM algorithm focused on log_2_ fold changes between replicates. Addressing missing data is beyond the scope of this study, which centers on computing GO term abundances across taxonomic levels in metaproteomics.

To further evaluate the accuracy of MiCId’s GO term abundance estimates across taxonomic levels, we analyzed the differences between MiCId’s reported log_2_ fold changes and the true values. Unipept was excluded from this analysis, as it does not compute GO term abundances across all identified taxa. The evaluation considered all possible pairwise comparisons (PWCs) between technical replicates: 9 PWCs from DFs 1–9 (UPS1 proteins spiked into yeast), 3 PWCs from DFs 10–12 (mixtures of 4 microorganisms), and 3 PWCs from DFs 19–21 (mixtures of 24 microorganisms). Accuracy was quantified using three metrics: expected error (E[Error]), expected mean absolute log_2_ fold change error (E[MALFCE]), and the percentage of errors with absolute values ≤ 2 compared to the true log_2_ fold changes (% EV). We define E[Error], E[MALFCE], and % EV as:

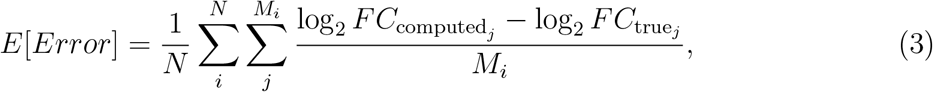

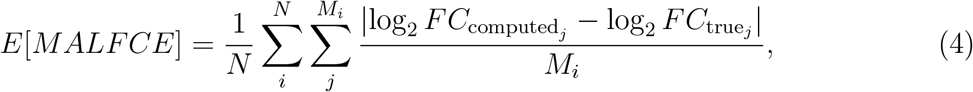

and

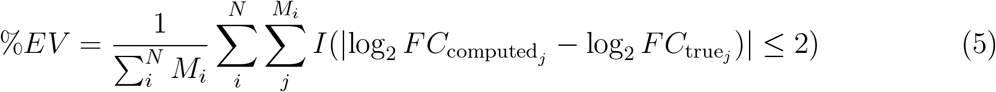

respectively. In equations 3–5, *N* denotes the total number of pairwise comparisons, and *M*_*i*_ is the number of log_2_ fold changes calculated in comparison *i*. As shown in Table 2, MiCId’s computed log_2_ fold changes for GO term abundances are consistently centered around zero across all taxonomic levels, as indicated by low E[Error] and E[MALFCE] values. Additionally, 92–98% of computed errors fall within ±2 of the true log_2_ fold change values (% EV), demonstrating the accuracy and precision of the EM algorithm. This threshold aligns with typical variability observed in quantitative proteomics.^58^

**Table 2:** Performance evaluation of MiCId’s computed GO term abundances across different taxonomic levels. To assess the accuracy of the GO term abundances reported by MiCId at various taxonomic levels, we computed the difference between the log_2_ fold changes of MiCId’s reported abundances and the true log_2_ fold changes for all possible pairwise comparisons (PWC) between technical replicates for data files DFs 1-9, DFs 10-12, and DFs 19-21. The table above shows the number of log_2_ fold change pairwise comparisons computed at each taxonomic level and used in the calculation of three metrics for evaluating MiCId’s computed GO term abundances: expected error (E[Error]), expected mean absolute log_2_ fold change error (E[MALFCE]), and the percentage of computed error values with an absolute value less than or equal to 2 compared to the true log_2_ fold change values (% EV).

| Results for data files 1-9 (48 proteins UPS1) |  |  |  |  |  |
| --- | --- | --- | --- | --- | --- |
| taxonomic Level | Nº PWC | Nº Go Terms | E[Error] | E[MALFCE] | % EV |
| Root | 9 | 8901 | 0.00 | 0.40 | 0.98 |
| Phylum | 9 | 9618 | 0.15 | 0.66 | 0.95 |
| Class | 9 | 9618 | 0.15 | 0.66 | 0.95 |
| Order | 9 | 9618 | 0.15 | 0.66 | 0.95 |
| Family | 9 | 9618 | 0.15 | 0.66 | 0.95 |
| Genus | 9 | 9618 | 0.15 | 0.66 | 0.95 |
| Species | 9 | 9618 | 0.15 | 0.66 | 0.95 |
| Results for data files 10-12 (Mixture of 4 microorganisms) |  |  |  |  |  |
| taxonomic Level | Nº PWC | Nº Go Terms | E[Error] | E[MALFCE] | % EV |
| Root | 3 | 1236 | 0.05 | 0.79 | 0.92 |
| Phylum | 3 | 1585 | 0.02 | 0.80 | 0.93 |
| Class | 3 | 1585 | 0.02 | 0.80 | 0.93 |
| Order | 3 | 2001 | 0.02 | 0.84 | 0.92 |
| Family | 3 | 2001 | 0.02 | 0.84 | 0.92 |
| Genus | 3 | 2001 | 0.01 | 0.84 | 0.92 |
| Species | 3 | 2001 | 0.02 | 0.84 | 0.92 |
| Results for data files 20-21 (Mixture of 24 microorganisms) |  |  |  |  |  |
| taxonomic Level | Nº PWC | Nº Go Terms | E[Error] | E[MALFCE] | % EV |
| Root | 3 | 1642 | -0.04 | 0.58 | 0.96 |
| Phylum | 3 | 3100 | 0.02 | 0.80 | 0.92 |
| Class | 3 | 4198 | 0.02 | 0.81 | 0.92 |
| Order | 3 | 5063 | 0.00 | 0.66 | 0.94 |
| Family | 3 | 5340 | 0.00 | 0.66 | 0.94 |
| Genus | 3 | 8300 | -0.01 | 0.64 | 0.95 |
| Species | 3 | 9128 | -0.02 | 0.62 | 0.95 |

Another key aspect highlighted in Table 2 is the number of GO terms identified and used for evaluating GO term abundances at each taxonomic level. For DFs 1-9, which consist of the *Homo sapiens* species, a total of 9,618 GO term abundances are computed for each taxonomic level. This consistent number of GO terms across all taxonomic levels is due to the single-species identification. For DFs 10-12, representing a mixture of four microorganisms, the number of GO term abundances varies by taxonomic level. Specifically, 1,595 GO terms are computed at the phylum and class levels, while 2,001 GO term abundances are computed at the order, family, genus, and species levels. The difference in the number of GO terms between the higher (phylum and class) and lower (order, family, genus, species) taxonomic levels arises from the distinct taxonomic lineages of the four identified species up to the order level. Beyond the order level, the number of unique lineages decreases, leaving only two distinct lineages. For DFs 20-21, which involve a mixture of 24 microorganisms, the number of GO term abundances ranges from 3,063 at the phylum level to 7,704 at the species level. This variation is due to the fact that each taxonomic level requires a different number of taxa to fully represent the identified species.

It is important to emphasize that, although the number of GO term abundances computed or identified may vary at different taxonomic levels, the number of unique GO terms remains constant across all levels for each data file. The observed differences in the number of GO terms are solely due to the varying number of taxa identified at different taxonomic level that share the same GO terms. These findings highlight the ability of the proposed EM algorithm to accurately compute biological function abundances across taxonomic levels, utilizing both shared and unshared peptides from confidently identified species. The GO term abundance values used to compute the results presented in Table 2 are provided in Supplementary Tables S95-S472.

### Analysis of human gut microbiome dataset

The human gut microbiome dataset used in this study was derived from stool samples collected from four children and was originally used for a deep metaproteomics analysis of the human gut microbiome.^9^ Among these samples, two were obtained from children diagnosed with ulcerative colitis (HM604, HM621), one sample came from a child with Crohn’s disease in remission (HM541), and one control sample was sourced from a child without IBD (HM609). In the original publication of this dataset, the MetaPro-IQ approach^59^ was used for protein/peptide identification, and the identified peptides were then used for taxonomic analysis with Unipept. We have now also used these identified peptides for functional analysis with Unipept (Supplementary Tables S494-S502). Therefore, in this manuscript, we refer to the results generated by MetaPro-IQ and Unipept as “MetaPro-IQ-Unipept”.

We compared genus level biomass compositions of human gut samples computed using the proposed EM algorithm (via Eq. S10) with those obtained from MetaPro-IQ-Unipept (via Eq. 1). The analysis showed a high degree of concordance, with a correlation coefficient of 0.96 between the average genus biomasses estimated by the two methods. Furthermore, PCA of species level biomass abundances and GO term molecular function abundances demonstrated the high reproducibility of the proposed EM algorithm. Technical replicates of the same sample clustered significantly closer together than replicates from different samples, indicating robust and consistent estimation of both taxonomic and functional biomass profiles (Supplementary Fig. S16). Similar PCA clustering patterns were observed across other taxonomic levels for both taxonomic and functional abundances. Abundance values computed for taxa and GO terms across taxonomic levels are provided in Supplementary Tables S487–S493.

In terms of protein identification, an average of 6,966 protein groups were detected per sample at the species level, corresponding to approximately 4,220 molecular functions, 2,593 biological processes, and 484 cellular components. Detailed results across taxonomic levels are provided in Supplementary Tables S473–S479. Although the number of GO terms reported varies by taxonomic level, the set of unique GO terms remains consistent, allowing for uniform attribution of functional contributions across taxa. This consistency highlights the effectiveness of the proposed EM algorithm in assigning biological function abundances—an advantage over the traditional LCA approach.

Table 3 summarizes the overlap of taxa identified by the MiCId workflow and those identified by MetaPro-IQ-Unipept. Across all samples, the average overlap is 97% ± 6% SD for phyla, 96% ± 7% SD for classes, 95% ± 6% standard deviation (SD) for orders, 88% ± 4% SD for families, and 90% ± 2% SD for genera. These results indicate that more than 88% of the taxa reported by MiCId are consistent with those identified using MetaPro-IQ-Unipept. Species level overlap was excluded from the analysis, as species nomenclature is more prone to variation across databases due to the use of synonymous names and due to the discovery of novel species in metagenomics deep sequencing, many of which do not yet have standardized taxonomic names in the NCBI taxonomy database.

**Table 3:**
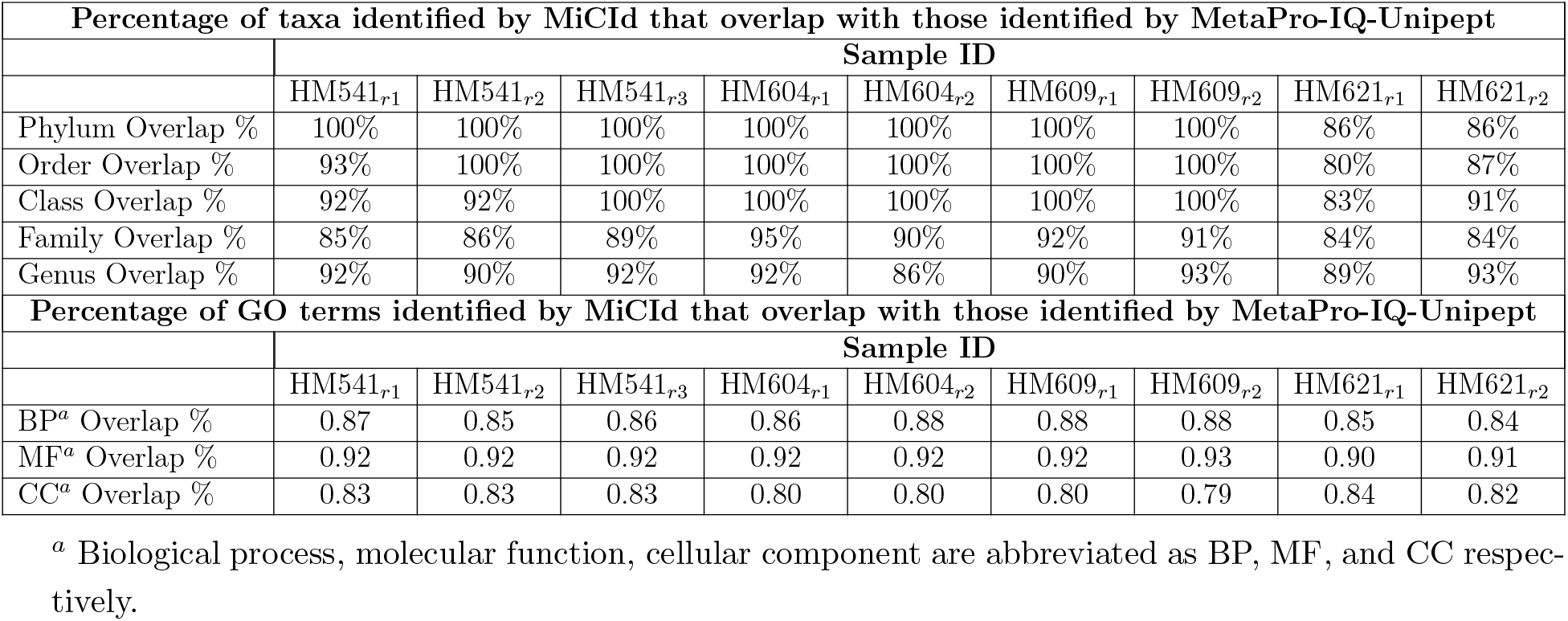
Percentage of taxa and GO terms identified by MiCId that overlap with those identified by MetaPro-IQ-Unipept for the human gut microbiome dataset.

| Percentage of taxa identified by MiCId that overlap with those identified by MetaPro-IQ-Unipept |  |  |  |  |  |  |  |  |  |
| --- | --- | --- | --- | --- | --- | --- | --- | --- | --- |
|  | Sample ID |  |  |  |  |  |  |  |  |
|  | HM541<sub>r1</sub> | HM541<sub>r2</sub> | HM541<sub>r3</sub> | HM604<sub>r1</sub> | HM604<sub>r2</sub> | HM609<sub>r1</sub> | HM609<sub>r2</sub> | HM621<sub>r1</sub> | HM621<sub>r2</sub> |
| Phylum Overlap % | 100% | 100% | 100% | 100% | 100% | 100% | 100% | 86% | 86% |
| Order Overlap % | 93% | 100% | 100% | 100% | 100% | 100% | 100% | 80% | 87% |
| Class Overlap % | 92% | 92% | 100% | 100% | 100% | 100% | 100% | 83% | 91% |
| Family Overlap % | 85% | 86% | 89% | 95% | 90% | 92% | 91% | 84% | 84% |
| Genus Overlap % | 92% | 90% | 92% | 92% | 86% | 90% | 93% | 89% | 93% |
| Percentage of GO terms identified by MiCId that overlap with those identified by MetaPro-IQ-Unipept |  |  |  |  |  |  |  |  |  |
|  | Sample ID |  |  |  |  |  |  |  |  |
|  | HM541<sub>r1</sub> | HM541<sub>r2</sub> | HM541<sub>r3</sub> | HM604<sub>r1</sub> | HM604<sub>r2</sub> | HM609<sub>r1</sub> | HM609<sub>r2</sub> | HM621<sub>r1</sub> | HM621<sub>r2</sub> |
| BP<sup>a</sup> Overlap % | 0.87 | 0.85 | 0.86 | 0.86 | 0.88 | 0.88 | 0.88 | 0.85 | 0.84 |
| MF<sup>a</sup> Overlap % | 0.92 | 0.92 | 0.92 | 0.92 | 0.92 | 0.92 | 0.93 | 0.90 | 0.91 |
| CC<sup>a</sup> Overlap % | 0.83 | 0.83 | 0.83 | 0.80 | 0.80 | 0.80 | 0.79 | 0.84 | 0.82 |

<sup>a</sup> Biological process, molecular function, cellular component are abbreviated as BP, MF, and CC respectively.

Additionally, Table 3 presents the percentage of GO terms identified by the MiCId workflow that were also detected by MetaPro-IQ-Unipept. Across all samples, the average overlap was 86% ± 2% SD for biological processes, 92% ± 1% SD for molecular functions, and 82% ± 2% SD for cellular components. This high degree of agreement in both taxonomic and functional annotations is particularly notable given that the underlying microbial databases were constructed from different sources. MetaPro-IQ employed a database based on the human gut microbial gene catalog,^60^ while MiCId utilized a database derived from human gut microbial genomes available in the NCBI BioSample database,^52^ as MiCId is currently designed to work directly with NCBI resources.

### Analysis of human oral microbiome dataset

The human oral microbiome dataset used consists of saliva and dental plaque samples from 12 children prone to dental caries. Each sample was incubated in biofilm reactors under two conditions—with sugar (WS) and without sugar (NS)—as described previously, ^45^ resulting in 24 samples. Metaproteomic data were originally processed using the Galaxy-P platform, with MEGAN5 used for taxonomic and functional analysis.

Table 4 quantifies the agreement between MiCId and methods from previous study by reporting the percentage of genera that were also identified by MEGAN5 and 16S rRNA analysis in the original study. On average, 82% of the genera identified by MiCId were also detected by MEGAN5, and 80% were detected by 16S rRNA. Further analysis of the 12 NS and 12 WS samples using 16S rRNA shows that, on average, 38.17 ± 4.24 SD genera were identified in NS samples, and 35.08 ± 3.20 SD in WS samples, based on genera with non-zero probe read counts. When focusing on genera with a relative biomass abundance of at least 1%, 16S rRNA identified an average of 6.33 ± 1.50 genera in NS samples and 4.17 ± 1.53 in WS samples. In contrast, MiCId identified an average of 14.25 ± 5.31 genera in NS samples and 6.33 ± 1.78 in WS samples overall. When considering only those genera with biomass abundance ≥1%, MiCId identified an average of 7.33 ± 1.67 genera for NS samples and 3.58 ± 1.00 for WS samples. Furthermore, the genus level biomass compositions across all NS and WS samples, as derived from MiCId, MEGAN5, and 16S rRNA, showed notable consistency in their average biomass profiles (Supplementary Fig. 17S).

**Table 4:**
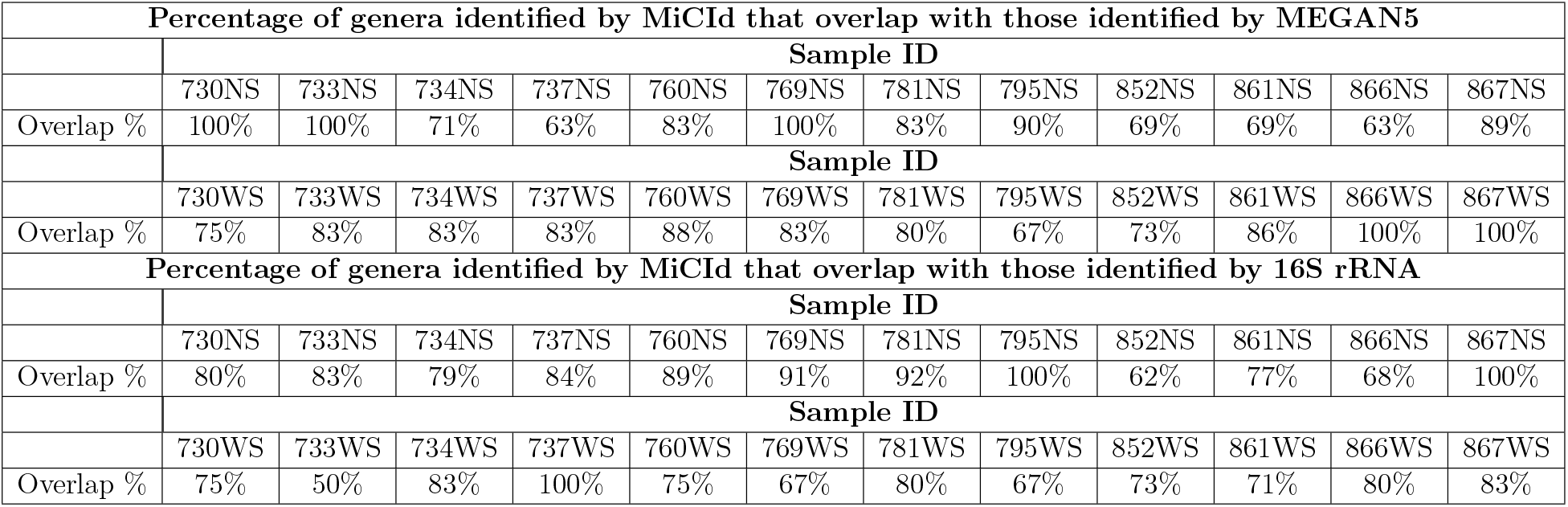
Percentage of genera identified by MiCId that overlap with those identified by MEGAN5 and 16S rRNA.

In parallel with the earlier analysis of this dataset, PCA was performed on the computed biomass abundances and GO term molecular function abundances at the species level. These values were derived using the proposed EM algorithm—species biomass abundances via Eq. S10 and functional abundances via Eq. S13. As detailed in the computational formalism of the modified EM algorithm (Supporting Information File), biological function abundances are calculated based on the computed conditional probability *p*(*k* | *t*_*α*_), representing the likelihood of biological function *k* originating from taxon *α*, as defined in Eq. S12. Panels (C) and (D) of Supplementary Fig. 17S display the PCA plots for species level biomass and molecular function abundances, respectively. The PCA results generated using the proposed EM algorithm are consistent with findings from the original study, showing a clear separation of NS and WS samples along the first principal component, except for samples 730NS and 733NS, which did not separate as distinctly.

At the species level, an average of 3,774 protein groups were identified in the NS samples, compared to 3,356 in the WS samples. These proteins were mapped to an average of 2,219 (NS) and 1,884 (WS) molecular function GO terms, 1,418 (NS) and 1,251 (WS) biological process GO terms, and 322 (NS) and 288 (WS) cellular component GO terms. Compared to the original study using MEGAN5,^45^ the proposed method identified, on average, 65% ± 3% (SD) of the GO terms across all 24 samples. The percentage of GO terms co-identified between MiCId and MEGAN5 falls within the range of values obtained in an evaluation of six metaproteomics software tools^22^ using oral microbiome samples from the same dataset. More importantly, a significant portion of the identified GO term molecular functions for the WS and NS samples in the current study were also identified and considered differentially expressed in the original study. Detailed information on the number of protein groups and associated biological functions at other taxonomic levels is provided in Supplementary Tables S473–S479.

It is important to emphasize that the proposed method leverages confidently identified proteins to infer GO terms, which are subsequently used by the EM algorithm to estimate their abundances across the taxonomic hierarchy of identified microorganisms. This approach ensures a consistent set of GO terms across taxonomic levels, with variations arising from the number of taxa associated with each GO term and their corresponding abundance estimates. These abundances differ both within and across taxonomic levels, depending on the number of taxa sharing a GO term and the extracted ion areas of peptides mapped to that term. Table 5 illustrates this by highlighting GO molecular functions that were elevated in WS compared to NS samples in the original study, ^45^ and similarly identified using the proposed method. The first half of the table presents expected GO term abundances at the genus level, while the second half provides species level abundances for selected terms. Notably, GO terms are shared among taxa across taxonomic levels—an outcome that cannot be achieved using the LCA algorithm. A complete list of identified GO terms, along with their computed abundances across taxonomic levels for each of the 24 samples, is provided in Supplementary Tables S480–S486.

**Table 5:** Average molecular function abundances for a few selected GO terms at both the genus and species levels from the human oral microbiome dataset, which consists of 12 samples incubated in biofilm reactors under two conditions: with sugar (WS) and without sugar (NS). Marked with a dot circle (•) are GO term molecular functions previously shown to be elevated in the WS samples in the original study. ^45^

| GO identifier | GO term description | Genus | Ab.<sup>a</sup> NS | Ab.<sup>a</sup> WS |
| --- | --- | --- | --- | --- |
| GO:0004345 | glucose-6-phosphate dehydrogenase | *Streptococcus* | 5.6E-06 | 3.3E-05 |
| GO:0004736 | pyruvate carboxylase | *Veillonella* | 4.7E-06 | 1.5E-05 |
| ●GO:0004356 | glutamine synthetase | *Streptococcus* | 1.8E-04 | 7.3E-04 |
| ●GO:0047112 | pyruvate oxidase | *Streptococcus* | 2.8E-05 | 5.8E-04 |
| GO:0004340 | glucokinase | *Streptococcus* | 4.4E-04 | 6.7E-04 |
| ●GO:0004084 | branched-chain-amino-acid transaminase | *Streptococcus* | 2.0E-05 | 2.6E-04 |
| GO:0004427 | diphosphate phosphatase | *Streptococcus* | 7.8E-04 | 1.9E-03 |
| GO:0004427 | diphosphate phosphatase | *Veillonella* | 1.3E-04 | 2.0E-04 |
| ●GO:0004459 | L-lactate dehydrogenase | *Streptococcus* | 1.1E-03 | 2.5E-03 |
| ●GO:0004459 | L-lactate dehydrogenase | *Granulicatella* | 9.8E-04 | 1.1E-03 |
| ●GO:0004634 | phosphopyruvate hydratase | *Veillonella* | 5.0E-05 | 3.1E-05 |
| ●GO:0004634 | phosphopyruvate hydratase | *Granulicatella* | 9.8E-05 | 2.5E-05 |
| ●GO:0004634 | phosphopyruvate hydratase | *Haemophilus* | 2.1E-04 | 3.3E-05 |
| ●GO:0004332 | fructose-bisphosphate aldolase | *Streptococcus* | 1.7E-03 | 3.8E-03 |
| ●GO:0004332 | fructose-bisphosphate aldolase | *Veillonella* | 9.7E-05 | 7.3E-05 |
| ●GO:0004332 | fructose-bisphosphate aldolase | *Granulicatella* | 7.0E-05 | 6.6E-05 |
| ●GO:0004332 | fructose-bisphosphate aldolase | *Haemophilus* | 1.6E-04 | 4.7E-05 |
| GO:0051287 | NAD binding | *Streptococcus* | 2.9E-03 | 9.1E-03 |
| GO:0051287 | NAD binding | *Veillonella* | 1.1E-04 | 1.6E-04 |
| GO:0004743 | pyruvate kinase | *Streptococcus* | 1.6E-04 | 5.5E-04 |
| GO:0004743 | pyruvate kinase | *Veillonella* | 7.8E-06 | 6.1E-06 |
| GO:0004743 | pyruvate kinase | *Haemophilus* | 4.6E-05 | 6.1E-06 |
| ●GO:0051536 | iron-sulfur cluster binding | *Streptococcus* | 4.6E-05 | 1.5E-04 |
| ●GO:0051536 | iron-sulfur cluster binding | *Veillonella* | 1.5E-04 | 2.3E-04 |
| GO:0004360 | glutamine-fructose-6-phosphate transaminase | *Streptococcus* | 1.3E-04 | 4.8E-04 |
| GO:0004360 | glutamine-fructose-6-phosphate transaminase | *Veillonella* | 6.8E-05 | 3.8E-05 |
| **Species level molecular function abundances for the above highlighted GO terms** |  |  |  |  |
| GO identifier | GO term description | Species | Ab. NS | Ab. WS |
| GO:0004345 | glucose-6-phosphate dehydrogenase | *Streptococcus gordonii* | 1.7E-06 | 6.1E-06 |
| GO:0004345 | glucose-6-phosphate dehydrogenase | *Streptococcus oralis* | 4.0E-07 | 5.2E-06 |
| GO:0004345 | glucose-6-phosphate dehydrogenase | *Streptococcus parasanguinis* | 5.6E-07 | 2.8E-06 |
| GO:0004345 | glucose-6-phosphate dehydrogenase | *Uncultured Streptococcus sp.* | 1.8E-07 | 1.8E-05 |
| ●GO:0051536 | iron-sulfur cluster binding | *Streptococcus gordonii* | 1.6E-05 | 1.4E-05 |
| ●GO:0051536 | iron-sulfur cluster binding | *Streptococcus oralis* | 1.1E-06 | 1.5E-05 |
| ●GO:0051536 | iron-sulfur cluster binding | *Streptococcus parasanguinis* | 2.0E-07 | 5.5E-06 |
| ●GO:0051536 | iron-sulfur cluster binding | *Uncultured Streptococcus sp.* | 2.5E-05 | 9.1E-05 |
| ●GO:0051536 | iron-sulfur cluster binding | *Uncultured Veillonella sp.* | 1.6E-05 | 5.6E-05 |
| ●GO:0051536 | iron-sulfur cluster binding | *Veillonella parvula* | 1.3E-04 | 7.0E-05 |
| GO:0004360 | glutamine-fructose-6-phosphate transaminase | *Streptococcus gordonii* | 3.5E-05 | 3.7E-05 |
| GO:0004360 | glutamine-fructose-6-phosphate transaminase | *Streptococcus oralis* | 3.9E-06 | 3.8E-05 |
| GO:0004360 | glutamine-fructose-6-phosphate transaminase | *Streptococcus parasanguinis* | 4.9E-06 | 4.5E-05 |
| GO:0004360 | glutamine-fructose-6-phosphate transaminase | *Uncultured Streptococcus sp.* | 1.0E-04 | 2.5E-04 |
| GO:0004360 | glutamine-fructose-6-phosphate transaminase | *Uncultured Veillonella sp.* | 4.6E-05 | 2.3E-05 |
| GO:0004360 | glutamine-fructose-6-phosphate transaminase | *Veillonella parvula* | 2.0E-05 | 1.1E-05 |

<sup>a</sup> Abundance is abbreviated as Ab.

As a specific example of how GO terms are shared and how their abundances can vary across taxonomic levels and samples, consider GO term GO:0004345 (highlighted in green in Table 5). At the genus level, this GO term is associated with *Streptococcus*, with computed abundances of 5.6E-06 (NS) and 3.3E-05 (WS). At the species level, it appears in four *Streptococcus* species: *Uncultured Streptococcus sp*. (1.8E-07 NS, 1.8E-05 WS), *S. oralis* (4.0E-07 NS, 5.2E-06 WS), *S. gordonii* (1.7E-06 NS, 6.1E-06 WS), and *S. parasanguinis* (5.6E-07 NS, 2.8E-06 WS). *S. gordonii* and *Uncultured Streptococcus sp*. are the main contributors in the NS and WS samples, respectively.

Similarly, GO term GO:0004360 (red in Table 5) is linked to both *Streptococcus* and *Veillonella*. At the species level, it appears in four *Streptococcus* and two *Veillonella species*. In the NS sample, the top contributors are *Uncultured Streptococcus sp*. (1.0E-04), *Uncultured Veillonella sp*. (4.6E-05), and *S. gordonii* (3.5E-05). In the WS sample, they are *Uncultured Streptococcus sp*. (2.5E-04), *S. parasanguinis* (4.5E-05), and *S. oralis* (3.8E-05).

### Advantages and Limitations

To further illustrate an advantage of MiCId’s EM algorithm over the commonly used LCA algorithm in identifying and quantifying microbial biological functions, we highlight results from data file 10 (DF-10), which are also included in Table 2. For this dataset, Unipept reports 342 unique biological process GO terms, but only 164 are correctly assigned at the species level. In contrast, MiCId reports 193 unique GO terms and successfully assigns them across all taxonomic levels, with 259 GO terms at the phylum and class levels, and 354 GO terms consistently assigned from order to species levels. While the total number of GO term assignments varies by taxonomic level due to differences in taxa, the set of unique GO terms remains consistent across all levels. This ability to compute GO term abundances across taxonomic levels for the same set of GO terms demonstrates a key advantage of MiCId’s EM algorithm over the LCA algorithm.

Another key advantage of assigning biological functions across the full taxonomic lineage is the ability to evaluate each taxon’s contribution to the community’s functional profile and to track how these contributions change under varying conditions. The EM algorithm further enhances this analysis by computing the abundance of a biological function *k* for taxon *α* (*t*_*α*_) as the joint probability *p*(*k*|*t*_*α*_)I:*p*(*t*_*α*_). These abundances are normalized such that their sum equals the biomass of *t*_*α*_ (i.e., ∑_*k*_ *p*(*k*|*t*_*α*_)*p*(*t*_*α*_) = *p*(*t*_*α*_)), making the functional analysis more intuitive. Consequently, the total abundance of GO terms associated with a biological pathway directly reflects a taxon’s contribution to that pathway.

In this study, we enhanced the MiCId workflow to enable identification and quantification of GO term abundances in metaproteomics. However, the workflow has certain limitations. Currently, MiCId supports only MS/MS data acquired in data-dependent acquisition (DDA) mode and DIA data with isolation windows of 2 Daltons or smaller. As a result, it cannot process DIA data from instruments like the TIMS-TOF mass spectrometer, which commonly uses larger isolation windows.^61^ In contrast, MiCId is compatible with DIA data from the Orbitrap Astral mass spectrometer, which typically uses 2 Dalton windows. ^62^ Another limitation is the lack of built-in differential expression analysis for GO term abundances. We plan to address these issues in future updates by extending DIA support and adding differential expression functionality.

Another current limitation of the MiCId workflow is its reliance on the NCBI database,^63^ which restricts analyses using protein and GO term databases derived from NCBI resources. Despite this constraint, NCBI remains a comprehensive and well-curated source of microbial data. For instance, there is an ongoing initiative at NCBI to curate and provide GO term annotations for bacterial proteins in the RefSeq database,^48^ and the BioSample database offers rich metadata and broad coverage of microorganisms from diverse environments.

In this study, we leveraged NCBI resources to construct representative databases for the human oral and gut microbiomes, and we showed that our results aligned well with findings from previous studies. Notably, MiCId allows users to create customized microorganismal databases by specifying NCBI taxonomic identifiers.^64^ To address current limitations in database construction, we will work in future versions of MiCId to support user-defined protein databases and provide an interface to interact with BioSample metadata, facilitating the creation of sample-specific databases. Additionally, although not yet implemented, the proposed EM algorithm is designed to be adaptable to other functional annotation sources beyond GO terms, such as the Enzyme Commission (EC),^65^ Kyoto Encyclopedia of Genes and Genomes (KEGG),^66^ and Clusters of Orthologous Genes (COG)^67^ databases.

## Conclusion

In conclusion, we have demonstrated that the newly augmented MiCId can accurately identify biological functions and that the integrated modified EM algorithm can estimate their abundances across the entire taxonomic lineage of confidently identified microorganisms in mass-spectrometry-based metaproteomics. The proposed EM algorithm provides a solution to the shared confidently identified peptide problem in biological function assignment across taxonomic levels, offering a novel way for assigning biological functions to the entire taxonomic lineage of confidently identified microorganisms. This ability of the proposed EM method contrasts with the commonly used LCA in metaproteomics. By accurately assigning biological functions across the entire taxonomic lineage of identified microorganisms, this approach provides an alternative means to study microbial community whose microorganismal protein databases can be derived from the NCBI database, enabling a more granular assessment of each taxon’s contribution to the biological functions within the community.

This work represents a novel method for biological function analysis in metaproteomics, and we expect that this new functionality in MiCId’s workflow will be beneficial to the massspectrometry-based metaproteomics community. To ensure accessibility, we have integrated the proposed approach into MiCId’s graphical user interface (GUI). Additionally, the source code for MiCId’s workflow, written in C++, is available for download along with the GUI. By making the source code publicly available, we aim to facilitate the integration of the proposed EM algorithm into other workflows. Detailed instructions on how to use the method are included in MiCId’s user manual. MiCId’s workflow, along with the source codes and executables, are freely available for the Linux environment and can be downloaded from https://www.ncbi.nlm.nih.gov/CBBresearch/Yu/downloads.html.

## AUTHOR INFORMATION

## Authors

Aleksey Y. Ogurtsov — Division of Intramural Research, National Library of Medicine, National Institutes of Health, Bethesda, MD 20894, USA;

## Author Contributions

G.A. and YK. Y. designed the study. G.A., A.O., and YK. Y. analyzed the data. G.A. and A.O. wrote computer programs to implement the proposed ideas of this study in MiCId’s workflow. G.A and YK. Y. prepared the manuscript. All authors have given approval to the final version of the manuscript

## Acknowledgment

We would like to thank Bart Mesuere for kindly provided us with the complete list of microorganisms included in Unipept. We thank the administrative group of the National Institutes of Health Biowulf Cluster where all the computational tasks were carried out.

## Supporting Information Available

Supporting Information: Supplementary tables (XLXS);

- Table S1: list of MS/MS data files used;
- Table S2: list of the scientific names and taxonomic identifiers for the organisms used to build MiCId’s and X!Tandem’s target databases;
- Table S3: list of false positive species identified for DFs 10-21;
- Tables S4-S24: GO term results for Unipept analysis;
- Tables S25-S66: GO term results for MetaGOmics analysis;
- Table S67: GO term gold standard;
- Tables S68-S94: GO term results for X!Tandem analysis;
- Tables S95-S493: GO term results for MiCId analysis;
- Tables S494-S502: GO term results for MetaPro-IQ with Unipept analysis. Supporting Information (PDF):
- Computational formalism details of the modified EM algorithm with biomass constraint;
- Protein clustering and unclustering procedures;
- Figures S1-S4: species biomass profiles;
- Figures S5-S13: assessment of GO term identification sensitivity and the proportion of false discoveries (PFD);
- Figures S14-S15: assessment of GO term abundances;
- Figures S16: biomass and PCA plots for the human gut microbiome dataset;
- Figures S17: biomass and PCA plots for the human oral microbiome dataset.

## TOC Graphic

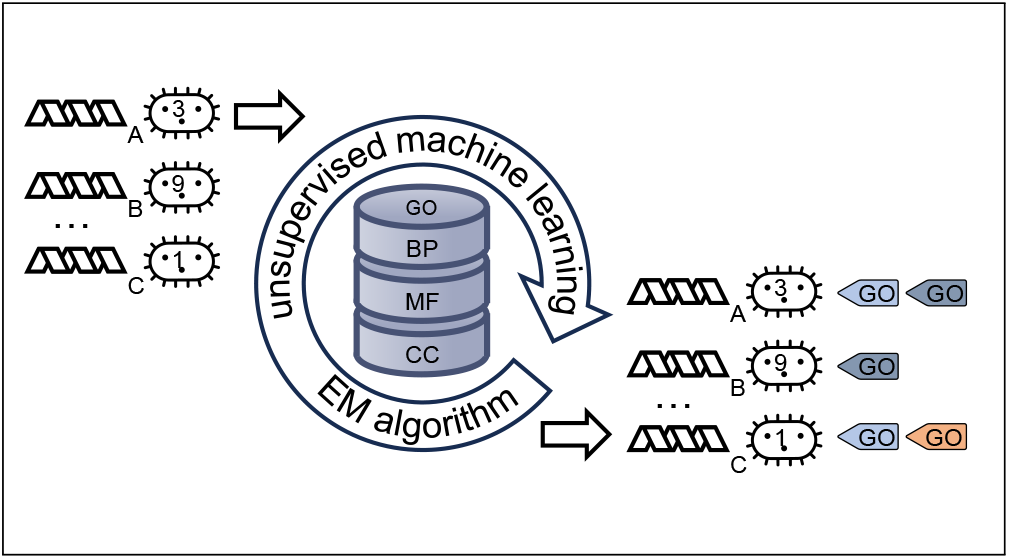

