## Supplementary File for "Biological Function Assignment Across Taxonomic Levels in Mass-Spectrometry-Based Metaproteomics via a Modified Expectation Maximization Algorithm"

### Table of contents


- Computational formalism details of the modified EM algorithm with biomass constraint;
- Protein clustering and unclustering procedures;
- Figures S1-S4: species biomass profiles;
- Figures S5-S13: assessment of GO term identification sensitivity and the proportion of false discoveries;
- Figures S14-S15: assessment of GO term abundances;
- Figures S16: biomass and PCA plots for the human gut microbiome dataset;
- Figures S17: biomass and PCA plots for the human oral microbiome dataset;
- Table S1 (XLSX): list of MS/MS data files used;
- Table S2 (XLSX): list of the scientific names and taxonomic identifiers for the organisms used to build MiCId's and X!Tandem's target databases;
- Table S3 (XLSX): list of false positive species identified for DFs 10-21;
- Tables S4-S24 (XLSX): GO term results for Unipept analysis;
- Tables S25-S66 (XLSX): GO term results for MetaGOmics analysis;
- Table S67 (XLSX): GO term gold standard;
- Tables S68-S94 (XLSX): GO term results for X!Tandem analysis;
- Tables S95-S493 (XLSX): GO term results for MiCId analysis;
- Tables S494-S502 (XLSX): GO term results for MetaPro-IQ with Unipept analysis.

### Computational formalism details of the modified EM algorithm with biomass constraint

The EM algorithm is a powerful statistical method used to maximize the likelihood of an observed outcome by adjusting model parameters and estimating the expectation value of hidden or unobserved variables. Since the distributions of these hidden variables depend on the model parameters, directly optimizing the likelihood becomes challenging. The EM algorithm simplifies this by breaking the process into three key steps: initialization, expectation, and maximization. In the first step, model parameters are initialized. In the second step, these parameters are used to compute the expected values of the hidden variables. Finally, in the third step, the model parameters are adjusted to maximize the likelihood using the expected values in place of the hidden variables. These steps are repeated iteratively, with the expectation and maximization steps performed recursively, until the algorithm converges to an optimal solution. For the problem at hand, the proposed EM algorithm is employed iteratively, once for each taxonomic level, to enable precise biological function analysis across different levels of microbial taxonomy.

We begin by establishing the required notation. In a metaproteomics experiment, thousands of peptides can be identified. Let $\Pi = \{\pi_1, \pi_2, \ldots, \pi_N\}$ represent a set of $N$ non-redundant peptides, each identified with an $E$-value $\leq 1$, and mapped to biological functions (i.e., GO terms) via confidently identified proteins. Assume that each $\pi_i$ has been observed with extracted ion chromatogram area, denoted by $n_i$. The $M$ taxa included at a specified taxonomy level are indexed by $\alpha$, where $t_\alpha$ represents the $\alpha$th taxon.

Let the variable $1 \leq k \leq K$ index the $K$ biological functions. Additionally, we define $d_\alpha$ as the peptidome of taxon $t_\alpha$. Let $p(\pi_{i,k} \leftarrow t_\alpha) \geq 0$ denote the probability that $\pi_i$, appearing in biological function $k$, originates from taxon $t_\alpha$. Let $0 \leq n_{i,k,\alpha} \leq n_{i,k}$ represent the extracted ion chromatogram area of $\pi_i$ in biological function $k$ coming from taxon $t_\alpha$. Clearly, the following conditions must hold: $\sum_\alpha n_{i,k,\alpha} = n_{i,k}$, and $\sum_{k=1}^{K} n_{i,k} = n_i$.

Since our application is not limited to using taxon-specific peptides, the taxon from which $\pi_i$ originates is not known in advance. Therefore, the values of $n_{i,k,\alpha}$ are treated as hidden variables. The objective of the EM algorithm is to estimate the parameters $p(\pi_{i,k} \leftarrow t_\alpha)$ that maximize the likelihood of the observed data $n_i$. Clearly, $\sum_\alpha p(\pi_{i,k} \leftarrow t_\alpha) I(\pi_i \in d_\alpha) = p(\pi_{i,k}) = \frac{n_{i,k}}{\sum_j n_{j,k}}$ where $p(\pi_{i,k})$ represents the prior probability that peptide $\pi_i$ is associated with biological function $k$, and the indicator function $I(\pi_i \in d_\alpha)$ equals 1 if $\pi_i$ belongs to the peptidome of $t_\alpha$ and 0 otherwise. As might be expected, we have that $\sum_{k=1}^{K} p(\pi_{i,k}) = p(\pi_i)$, which is the prior probability of peptide $\pi_i$, and $\sum_{i=1}^{N} p(\pi_i) = 1$.

We now describe the formal EM procedure, followed by our modification and simplification. Assuming independence among peptides' occurrences, we may write the likelihood for $\{n_{i,k,\alpha}\}$ consistent with the observed $\{n_{i,k}\}$ (assuming that $p(\pi_{i,k} \leftarrow t_\alpha)$s are known) as

$$L\left(\{n_{i,k,\alpha}\} | \{p(\pi_{i,k} \leftarrow t_\alpha)\}\right) = \frac{\left(\sum_{i=1}^{N} \sum_{k=1}^{K} n_{i,k}\right)!}{\prod_{i'=1}^{N} \prod_{k'=1}^{K} \prod_{\alpha=1}^{M} n_{i',k',\alpha}!} \prod_{i'=1}^{N} \prod_{k'=1}^{K} \prod_{\alpha=1}^{M} [p(\pi_{i',k'} \leftarrow t_\alpha)]^{n_{i',k',\alpha}}. \quad (1)$$

The first step of EM is to compute the expected values of the hidden variables. Because $n_{i,k} = \sum_\alpha n_{i,k,\alpha} I(\pi_i \in d_\alpha)$, the expected value of $n_{i,k,\alpha}$ for a fixed $i$ can be easily computed for any $t_\alpha$ whose peptidome contain $\pi_i$. Let $\varsigma(i,k)$ denote a realization of $\{n_{i,k,\alpha}\}$ satisfying $n_{i,k} = \sum_\alpha n_{i,k,\alpha} I(\pi_i \in d_\alpha)$. The expected value of $n_{i,k,\beta}$ can be written as

$$\begin{aligned} E[n_{i,k,\beta}|\{p(\pi_{j,k}\leftarrow t_\alpha)\}] &= \frac{\sum_{\varsigma(i,k)} L\left(\{n_{i,k,\alpha}\}|\{p(\pi_{j,k}\leftarrow t_\alpha)\}\right) n_{i,k,\beta}}{\sum_{\varsigma(i,k)} L\left(\{n_{i,k,\alpha}\}|\{p(\pi_{j,k}\leftarrow t_\alpha)\}\right)} \\ &= \frac{\partial \ln[\sum_{\varsigma(i,k)} L\left(\{n_{i,k,\alpha}\}|\{p(\pi_{j,k}\leftarrow t_\alpha)\}\right)]}{\partial \ln[p(\pi_{i,k}\leftarrow t_\beta)]} \end{aligned} \tag{2}$$

Because

$$\sum_{\varsigma(i,k)} L\left(\{n_{i,k,\alpha}\}|\{p(\pi_{j,k}\leftarrow t_\alpha)\}\right) \propto \left[\sum_\alpha p(\pi_{i,k}\leftarrow t_\alpha) I(\pi_i \in d_\alpha)\right]^{n_{i,k}}$$

we find

$$E[n_{i,k,\beta}|\{p(\pi_{j,k}\leftarrow t_\alpha)\}] = n_{i,k} \frac{p(\pi_{i,k}\leftarrow t_\beta) I(\pi_i \in d_\beta)}{\sum_\alpha p(\pi_{i,k}\leftarrow t_\alpha) I(\pi_i \in d_\alpha)} . \tag{3}$$

In the second step (maximization), one finds the next set of $\{p^{(\ell+1)}(\pi_{i,k} \leftarrow t_\alpha)\}$ to replace the previous set of $\{p^{(\ell)}(\pi_{i,k} \leftarrow t_\alpha)\}$ that was used to compute the expected values $E[n_{i,k,\beta}|\{p^{(\ell)}(\pi_{j,k} \leftarrow t_\alpha)\}]$. This is done by maximizing the log likelihood in Eq. (1) (with the $n_{i,k,\alpha}$s replaced by their expected values and treated as known numbers) via differentiation with respect to $\{p(\pi_{i,k}\leftarrow t_\alpha)\}$. There is of course a constraint that $1 = \sum_{i=1}^N p(\pi_i) = \sum_{i=1}^N \sum_{\alpha=1}^M p(\pi_i \leftarrow t_\alpha) = \sum_{i=1}^N \sum_{k=1}^K \sum_{\alpha=1}^M p(\pi_{i,k} \leftarrow t_\alpha)$. Introducing this constraint via a Lagrange multiplier, one ends up having

$$p^{(\ell+1)}(\pi_{i,k}\leftarrow t_\alpha) = \frac{E[n_{i,k,\alpha}|\{p^{(\ell)}(\pi_{j,k}\leftarrow t_\beta)\}]}{\sum_{i=1}^N \sum_{k=1}^K \sum_{\alpha=1}^M E[n_{i,k,\alpha}|\{p^{(\ell)}(\pi_{j,k}\leftarrow t_\beta)\}]} . \tag{4}$$

A few considerations lead us to take a simplified form of the above EM procedure. First, it requires a lot of data to faithfully estimate $\{p(\pi_{i,k}\leftarrow t_\alpha)\}$. Therefore, we make the simple choice that

$$p(\pi_{i,k}\leftarrow t_\alpha) \Rightarrow p(\pi_i\leftarrow t_\alpha) p(k|t_\alpha) \Rightarrow p(t_\alpha) p(\pi_i) p(k|t_\alpha) \tag{5}$$

to reduce the number of parameters to be fitted. Note

$$\sum_{k=1}^K p(k|t_\alpha) = 1.$$

Second, even though peptide $\pi_i$ is identified, there is no good way to infer its total ion count $n_i$ as there might be multiple identifications (some more confident and some less confident) or unfragmented $MS^1$ ions of the same peptide $\pi_i$. To simplify, we retain only the most confident identification per peptide weighted by the identification confidence

$$Z[E(\pi_i)] = \frac{n_i}{1 + E(\pi_i)/E_c} \equiv z_i. \tag{6}$$

Basically, we will replace $n_i$ by $z_i$ and $n_{i,k,\alpha}$ ($n_{i,k}$) by $z_{i,k,\alpha}$ ($z_{i,k}$). Here $E_c$ is the $E$-value cutoff used to control the expected number of FPs. The expected number of FP peptides identified is strongly controlled to be no more than 100 by setting $E_c$ equal to 100 divided by the total number of MS/MS spectra.

With the simplifications and modifications above, Eq. (3) becomes

$$\begin{aligned} E[z_{i,k,\beta}|\{p(t_\alpha), p(k|t_\alpha)\}] &= z_{i,k}\frac{p(t_\beta)p(\pi_i)p(k|t_\beta)I(\pi_i \in d_\beta)}{\sum_\alpha p(t_\alpha)p(\pi_i)p(k|t_\alpha)I(\pi_i \in d_\alpha)} \\ &= z_{i,k}\frac{p(t_\beta)p(k|t_\beta)I(\pi_i \in d_\beta)}{\sum_\alpha p(t_\alpha)p(k|t_\alpha)I(\pi_i \in d_\alpha)}\,. \end{aligned} \tag{7}$$

Before proceeding to maximization, we first note that the number of variables has decreased substantially under the simplification.

Here, we present a simplified form of the likelihood function in Eq. (1). Assuming independence among the occurrences of identified peptides, we can express the simplified likelihood function for the hidden variables $n_{i,k,\alpha}$, consistent with the observed data $n_{i,k}$, as follows

$$\begin{aligned} &L\left(\{z_{i,k,\alpha}\}|\{p(\pi_{i,k}\leftarrow t_\alpha)\}\right) \Rightarrow G(z_{i,k,\alpha})\prod_{i=1}^{N}\prod_{k=1}^{K}\prod_{\alpha=1}^{M}\left[p(\pi_i)p(t_\alpha)p(k|t_\alpha)\right]^{z_{i,k,\alpha}} \\ &= G(z_{i,k,\alpha})\left\{\prod_{i=1}^{N}\left[p(\pi_i)\right]^{z_i}\right\}\left\{\prod_{\alpha=1}^{M}\left[p(t_\alpha)\right]^{\sum_i z_{i,\alpha}}\right\}\left\{\prod_{k=1}^{K}\prod_{\alpha=1}^{M}\left[p(k|t_\alpha)\right]^{\sum_i z_{i,k,\alpha}}\right\} \\ &\equiv G(z_{i,k,\alpha})\cdot L_0(\{z_i\}|\{p(\pi_i)\})\cdot L_1(\{z_{i,\alpha}\}|\{p(t_\alpha)\})\cdot L_2(\{z_{i,k,\alpha}\}|\{p(k|t_\alpha)\}), \end{aligned} \tag{8}$$

where $G(z_{i,k,\alpha})$ is given by

$$G(z_{i,k,\alpha}) = \frac{\Gamma(1+\sum_i\sum_k z_{i,k})}{\prod_i\prod_k\prod_\alpha\Gamma(z_{i,k,\alpha}+1)}.$$

In our implementation of the EM algorithm with constraints, we first maximize the likelihood function $L_1$, subject to the constraint $\sum_\alpha p(t_\alpha) = 1$, to obtain the recursion for $p(t_\alpha)$ until convergence. During this process, the prior $p(\pi_i)$ for each $\pi_i$ is initially set to $1/N$. Setting

$$0 = \frac{d\ln L_1}{d\,p(t_\alpha)}$$

gives

$$\begin{aligned} p^{(\ell+1)}(t_\alpha) &\propto \sum_{i=1}^{N} E[z_{i,\alpha}|\{p(t_\alpha)\}] \\ &= \sum_{i=1}^{N} z_i\frac{p(t_\alpha)I(\pi_i \in d_\alpha)}{\sum_\beta p(t_\beta)I(\pi_i \in d_\beta)} \equiv C_\alpha^{(\ell+1)}. \end{aligned} \tag{9}$$

Along with the constraint condition $\sum_\alpha p(t_\alpha) = 1$, this leads to

$$p^{(\ell+1)}(t_\beta) \Rightarrow \frac{C_\beta^{(\ell+1)}}{\sum_\alpha C_\alpha^{(\ell+1)}} \,. \tag{10}$$

Once the values of $p(t_\alpha)$'s have converged in Eq. 9, we then proceed to maximize $L_2$ with respect to $p(k|t_\alpha)$, subject to the constraint that $\sum_k p(k|t_\alpha) = 1 \; \forall \; \alpha$. Setting

$$0 = \frac{d \ln L_2}{d\, p(k|t_\alpha)}$$

leads to

$$\begin{aligned} p^{(\ell+1)}(k|t_\alpha) &\propto \sum_{i=1}^{N} E[z_{i,k,\alpha}|\{p(t_\alpha),\; p^{(\ell)}(k|t_\alpha)\}] \\ &= \sum_{i=1}^{N} z_{i,k} \frac{p(t_\alpha)p^{(\ell)}(k|t_\alpha)I(\pi_i \in d_\alpha)}{\sum_\beta p(t_\beta)p^{(\ell)}(k|t_\beta)I(\pi_i \in d_\beta)} \equiv C_{k,\alpha}^{(\ell+1)}, \end{aligned} \tag{11}$$

The constraints conditions $\sum_k p(k|t_\alpha) = 1$ lead to

$$p^{(\ell+1)}(r|t_\beta) \Rightarrow \frac{C_{r,\beta}^{(\ell+1)}}{\sum_k C_{k,\alpha}^{(\ell+1)}} \,. \tag{12}$$

In the first iteration of the EM algorithm for the $L_1$ part, the priors $p(t_\alpha)$ are initialized to $1/M$, where $M$ is the total number of identified taxa. The expectation and maximization steps are repeated until numerical convergence is achieved for the priors $p(t_\alpha)$. Once the taxon priors are determined, the algorithm proceeds to the $L_2$ EM step. Here, the $p(k|t_\alpha)$ values are initialized to $1/K$, where $K$ is the total number of GO terms. After the first iteration of expectation and maximization, the values obtained for $p(k|t_\alpha)$ are used to compute GO term abundances (GA) for each identified taxon $\alpha$ ($t_\alpha$) as:

$$GA(k|t_\alpha) = p(k|t_\alpha)p(t_\alpha). \tag{13}$$

After numerical convergence is achieved for the $p(k|t_\alpha)$s values, these are used to compute enriched GO term abundances via Eq. 13. As has been shown,[1] the estimated expected probability $p(t_\alpha)$ can be interpreted as the relative biomass abundance of each identified $t_\alpha$, while the estimated $p(k|t_\alpha)$ represents the relative biomass abundance from identified $t_\alpha$ in biological function $k$. It is also important to mention that although we have used in our computational formalism the extracted ion chromatogram areas for the values of $\{n_i\}$,[2] these values can be substituted with any other quantity suitable to estimate relative biomass abundance such as spectral counting.[3,4]

There is a wealth of literature on the applications of the EM algorithm to solve computational problems in bioinformatics; here are a few useful references for the readers.[5–8] Additionally, we would like to emphasize that implementing the proposed EM algorithm requires only the use of the equations 9-12.

### Protein clustering and unclustering procedures

In our proposed method, confidently identified proteins serve as the basis for mapping identified peptides to GO terms. Since the statistical framework used for protein identification and its application have been detailed in our previous publications,[9,10] we now focus on describing the new clustering and unclustering procedures for protein identification that have been integrated into MiCId's workflow. We begin by outlining the protein clustering procedure, followed by the protein unclustering procedure. Additionally, an algorithm detailing the clustering procedure is provided in this subsection.

##### Protein clustering procedure

Our clustering strategy follows an iterative and transitive approach, with details provided in Algorithm S1 and described below:
1. **Sorting of Identified Proteins:**
First, all identified proteins are sorted in descending order based on the number of non-redundant confidently identified peptides, each with an $E$-value $\leq 1$. For proteins that share the same set of identified peptides, the order in which they appear in the sorted list does not matter, as they will be clustered together during the clustering procedure. Additionally, for proteins with an equal number of identified peptides, the order in which they appear in the sorted list is not particularly relevant, as they are likely to be clustered together if they share 50% or more of their weighted peptides during the clustering procedure; otherwise, they will not be clustered together.
2. **Weight Assignment to Non-Redundant Confidently Identified Peptides:**
For each non-redundant confidently identified peptide $\pi_i$, a weight $w(\pi_i)$ is assigned based on the peptide's $E$-value $E(\pi_i)$. This weight is used to compute the fraction of peptides shared between protein clusters. The weight $w(\pi_i)$ is calculated using the following equation:

$$w(\pi_i) = \frac{1}{1 + \frac{E(\pi_i)}{E_{cutoff}}} , \tag{14}$$

where $E_{cutoff}$ is the $E$-value used to control the proportion of false discoveries at 5%. Note that $w(\pi_i)$ values are bounded between zero and one. The fraction of peptides shared between protein $P_i$, which has $n_i$ non-redundant identified peptides, and protein $P_j$, which has $n_j$ non-redundant identified peptides, with $n_i \geq n_j$, is given by:

$$\rho(P_i, P_j) = \frac{\sum_k^{n_j} w(\pi_{jk}) I(\pi_{jk} \in P_i)}{\sum_k^{n_j} w(\pi_{jk}).} \tag{15}$$

In the above equation, $\pi_{jk}$ represents the $k^{th}$ peptide that belongs to species $j$, and $I(\pi_{jk} \in P_i)$ is an indicator function, which takes the value of one if peptide $\pi_{jk}$ also belongs to $P_i$, and zero otherwise.
3. **Protein Cluster Merge Condition:**

The protein cluster merge condition parameter $\Omega$ is initialized to 100%. Two protein clusters are merged when the fraction of weighted peptides shared between two clusters is greater than or equal to $\Omega$.
4. **Iterative Clustering:**
The clustering procedure begins with the first protein in the sorted list of proteins as the reference protein. All other lower-ranking proteins that share at least $\Omega$ of weighted non-redundant confidently identified peptides with the reference protein will have their cluster indexes updated to match the reference protein's cluster. The reference point is then moved to the next protein in the sorted list, and the process repeats. This continues until the reference point moves through all proteins in the list.
5. **Adjustment of the Merge Condition:**
After completing the clustering iteration with the current value of $\Omega$, the merge condition parameter $\Omega$ is decreased by 2.5%. While the value of $\Omega$ is $\geq 50\%$ the fourth step is repeated.
6. **Final Cluster Assignment:**
In the final step, the protein with the most significant $E$-value within a protein cluster (containing one or more proteins) is designated as the head of that cluster. The remaining proteins within that cluster are considered its members.

An exception to the aforementioned clustering rule is introduced to appropriately emphasize a protein cluster's unique evidence peptides—peptides that are not shared by other protein clusters. These unique peptides are critical for distinguishing one protein cluster from another. A protein is considered to have unique evidence peptides if it possesses two peptides, both with an $E$-value less than 10 divided by the total number of MS/MS spectra analyzed. Under this condition, the clustering procedure will not merge this protein cluster with others. The reasoning behind using 10 divided by the total number of MS/MS spectra analyzed as the $E$-value threshold ($E_t$) for determining whether a peptide is unique to a protein cluster is as follows: at this threshold, assuming accurate $E$-values (as demonstrated in previous studies[11]), there should be no more than 10 false positives identified peptides. This threshold is designed to control the emergence of false-positive protein clusters by minimizing the likelihood of incorrectly merging distinct clusters based on shared peptide evidence.

#### Protein unclustering procedure

The first step in the protein unclustering procedure is to select a set of confidently identified protein clusters. This is done by choosing protein clusters generated during the clustering procedure that pass the 1% FDR threshold and have $E$-values $\leq 1$. The next step involves separating protein members within a cluster that belong to species different from the species of the protein head of the cluster. In cases where a protein cluster contains multiple members from the same species, only the protein with the best $E$-value is selected to form a separate protein cluster. In the case of a tie in the best $E$-value for proteins, we opt to be conservative and do not select any of the proteins.

The unclustering procedure is repeated for each taxonomic level, and non-redundant protein records (WPs) for the protein head of the clusters are then used to retrieve GO terms from MiCId's biological function database. Non-redundant confidently identified peptides from these proteins are mapped to the corresponding GO terms. If a WP is used to query MiCId's biological function database and no GO term entry is available for that query, MiCId returns GO:XXXXXXX as the GO term. An additional modification involves introducing a root taxonomic level. At this level, the protein heads of the clusters are used—without applying the unclustering procedure—to obtain GO terms at the root level, which represents the highest possible taxonomic level. At the root level, no taxonomic differentiation is made for the identified GO terms.

**Algorithm 1** Protein Clustering Procedure

**Require:** $([\pi_1,\pi_2,\ldots,\pi_M])$ ▷ List of non-redundant confidently identified peptides.
**Require:** $([P_1,P_2,\ldots,P_N])$ ▷ List of identified proteins.
**Require:** $([E(\pi_1),\mathrm{E}(\pi_2),\ldots,\mathrm{E}(\pi_M)])$ ▷ List of non-redundant confidently identified peptides $E$-values.
**Require:** $([n_u(P_1),n_u(P_2),\ldots,n_u(P_N)])$ ▷ List storing the number of unique confidently peptides identified with $E$-value $\leq E_t$ mapped to $P_i$.
**Require:** $E_{cutoff}$ ▷ $E_{cutoff}$ is the $E$-value used to control the peptide proportion of false discoveries at 5%.

Set $\Omega = 1$
Set $n_u = 2$ ▷ Minimum number of unique confidently identified peptide that a protein $P_j$ must have in order for it not to cluster.
Set $\mathcal{C} = [\emptyset]$ ▷ Store the indices of proteins that have been clustered.
SORT($[P_1,P_2,P_3,...,P_N]$) ▷ Sort protein in decreasing order based on the number of non-redundant identified peptides mapped to protein $P_i$.
**for** $i = 1,2,\ldots,M$ **do** ▷ Computing weights to non-redundant confidently identified peptides.
    $w(\pi_i) = \frac{1}{1+\frac{E(\pi_i)}{E_{cutoff}}}$
**end for**
**while** $\Omega \geq 0.50$ **do**
    **for** $i = 1,2,\ldots,N$ **do**
        **if** $i \notin \mathcal{C}$ **then**
            **for** $j = i+1,\ldots,N$ **do**
                **if** $j \notin \mathcal{C}$ **then**
                    $\rho(P_i,P_j) = \frac{\sum_k^{n_j} w(\pi_{jk}) I(\pi_{jk} \in P_i)}{\sum_k^{n_j} w(\pi_{jk})}$
                    **if** $\rho(P_i,P_j) \geq \Omega$ and $n_u(P_j) < n_u$ **then** ▷ Condition for clustering.
                        add $j$ to $\mathcal{C}$ ▷ Protein $j$ has clustered to protein $i$ and the index $j$ is added to the list of clustered proteins $\mathcal{C}$.
                    **end if**
                **end if**
            **end for**
        **end if**
    **end for**
    $\Omega = \Omega - 0.025$ ▷ Updating the value of $\Omega$.
**end while**

### Figure S1

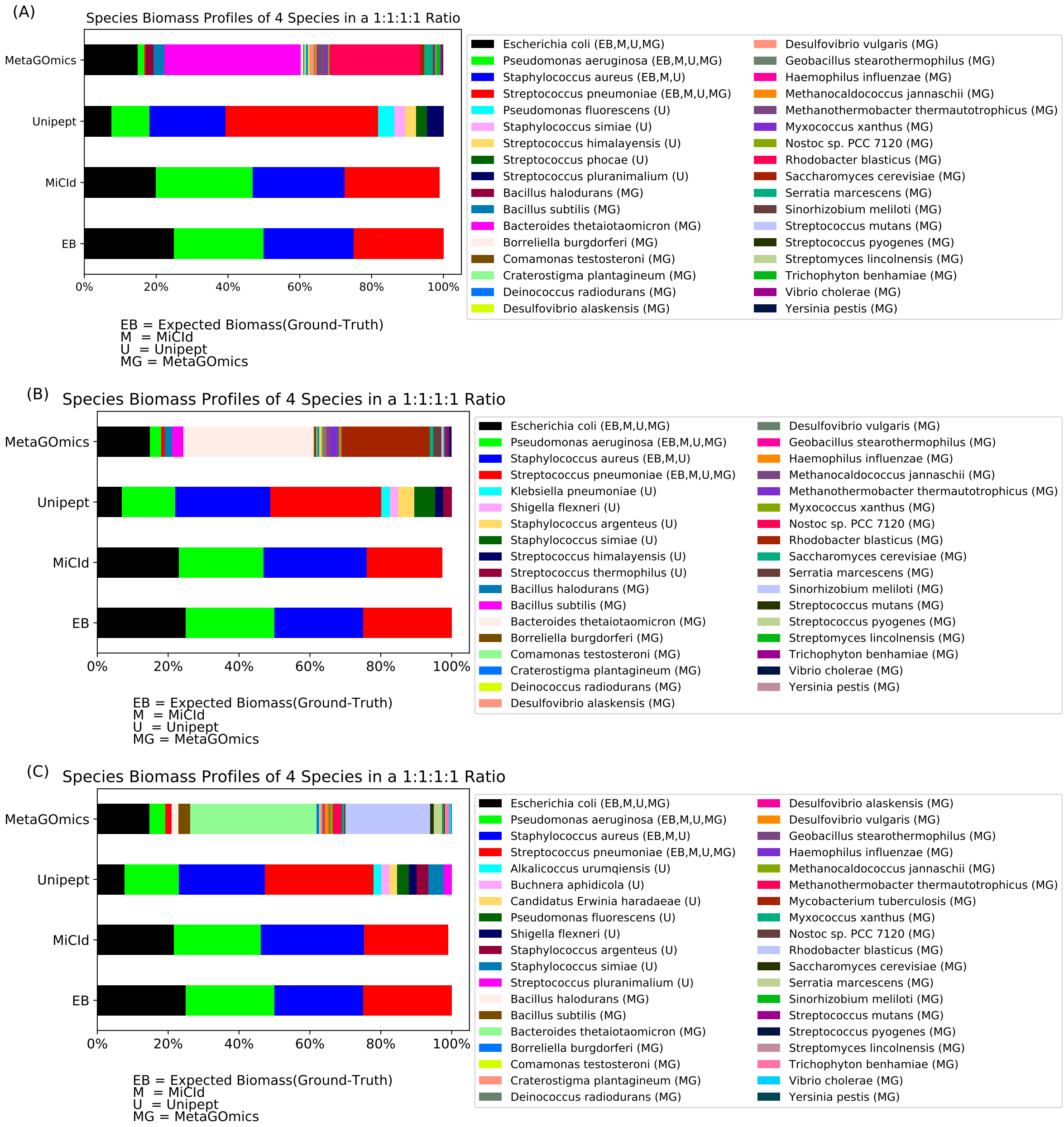


Figure S1: Comparison of species level biomass profiles reported MiCId and Unipept. Panels (A)-(C) present a comparison of species level biomass profiles, displayed as stacked bar plots showing species biomass computed by MiCId, Unipept, MetaGomics, and the expected biomass values (EB). Each stacked bar plot is accompanied by a list of the species included, with true positive species listed first, followed by any false positives identified. Panel (A)-(C) shows stacked bar plots for a sample composed of Staphylococcus aureus, Pseudomonas aeruginosa, Escherichia coli, and Streptococcus pneumoniae in a 1:1:1:1 ratio (data file 10-12).

### Figure S2

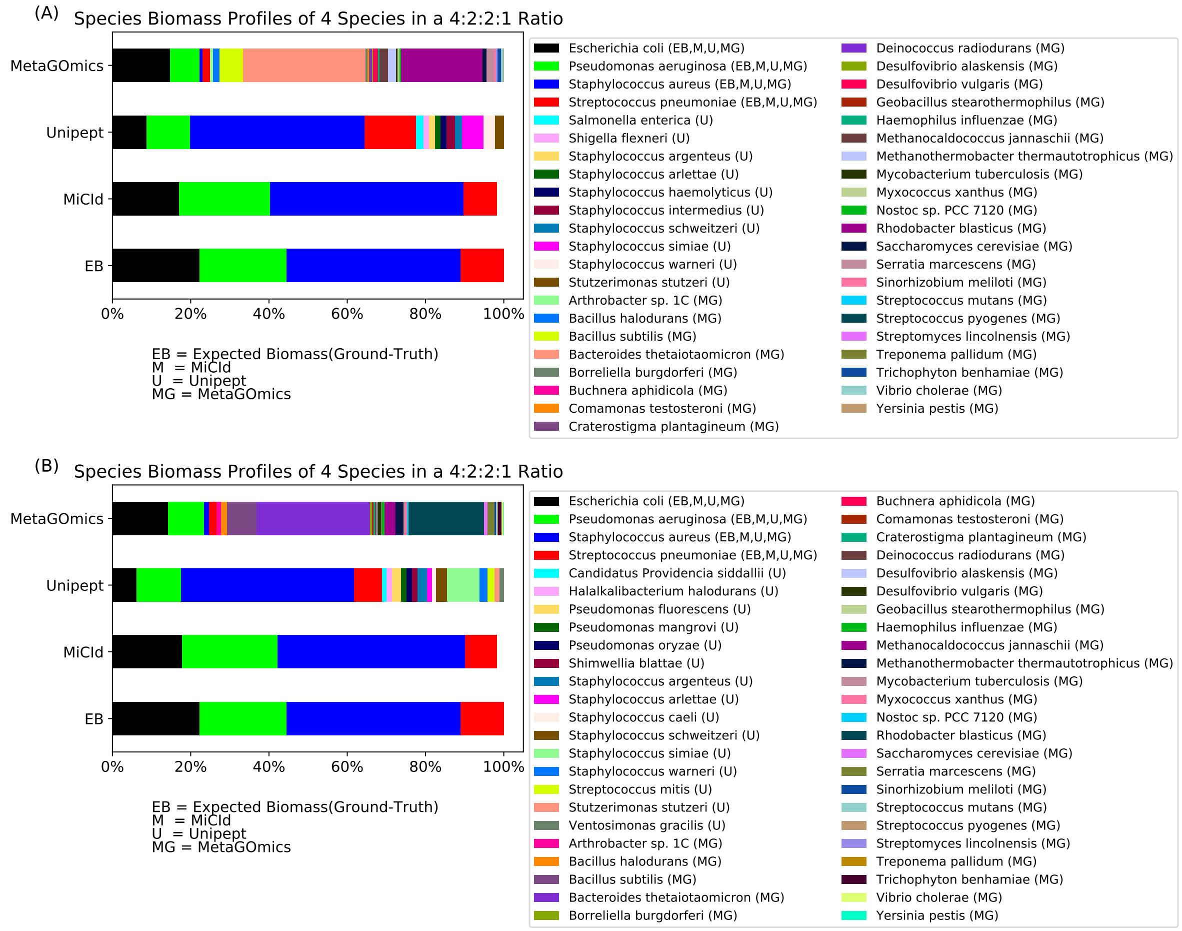


Figure S2: Comparison of species level biomass profiles reported MiCId and Unipept. Panels (A)-(B) present a comparison of species level biomass profiles, displayed as stacked bar plots showing species biomass computed by MiCId, Unipept, MetaGomics, and the expected biomass values (EB). Each stacked bar plot is accompanied by a list of the species included, with true positive species listed first, followed by any false positives identified. Panel (A)-(B) shows stacked bar plots for a sample composed of Staphylococcus aureus, Pseudomonas aeruginosa, Escherichia coli, and Streptococcus pneumoniae in a 4:2:2:1 ratio (data file 14-15)

### Figure S3

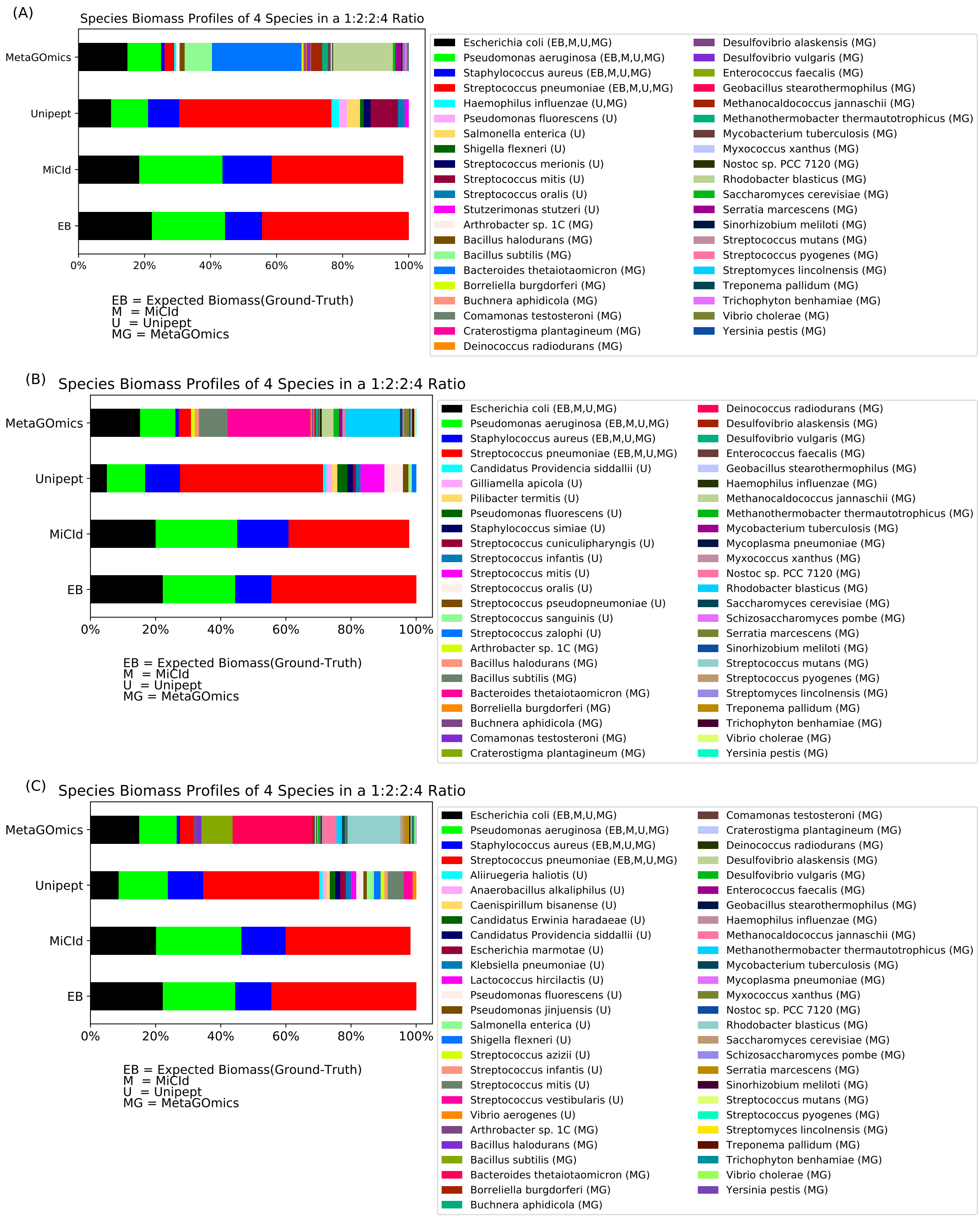


Figure S3: Comparison of species level biomass profiles reported MiCId and Unipept. Panels (A)-(C) present a comparison of species level biomass profiles for a sample composed of Staphylococcus aureus, Pseudomonas aeruginosa, Escherichia coli, and Streptococcus pneumoniae in a 1:2:2:4 ratio (data file 16-18), displayed as stacked bar plots showing species biomass computed by MiCId, Unipept, MetaGomics, and the expected biomass values (EB).

### Figure S4

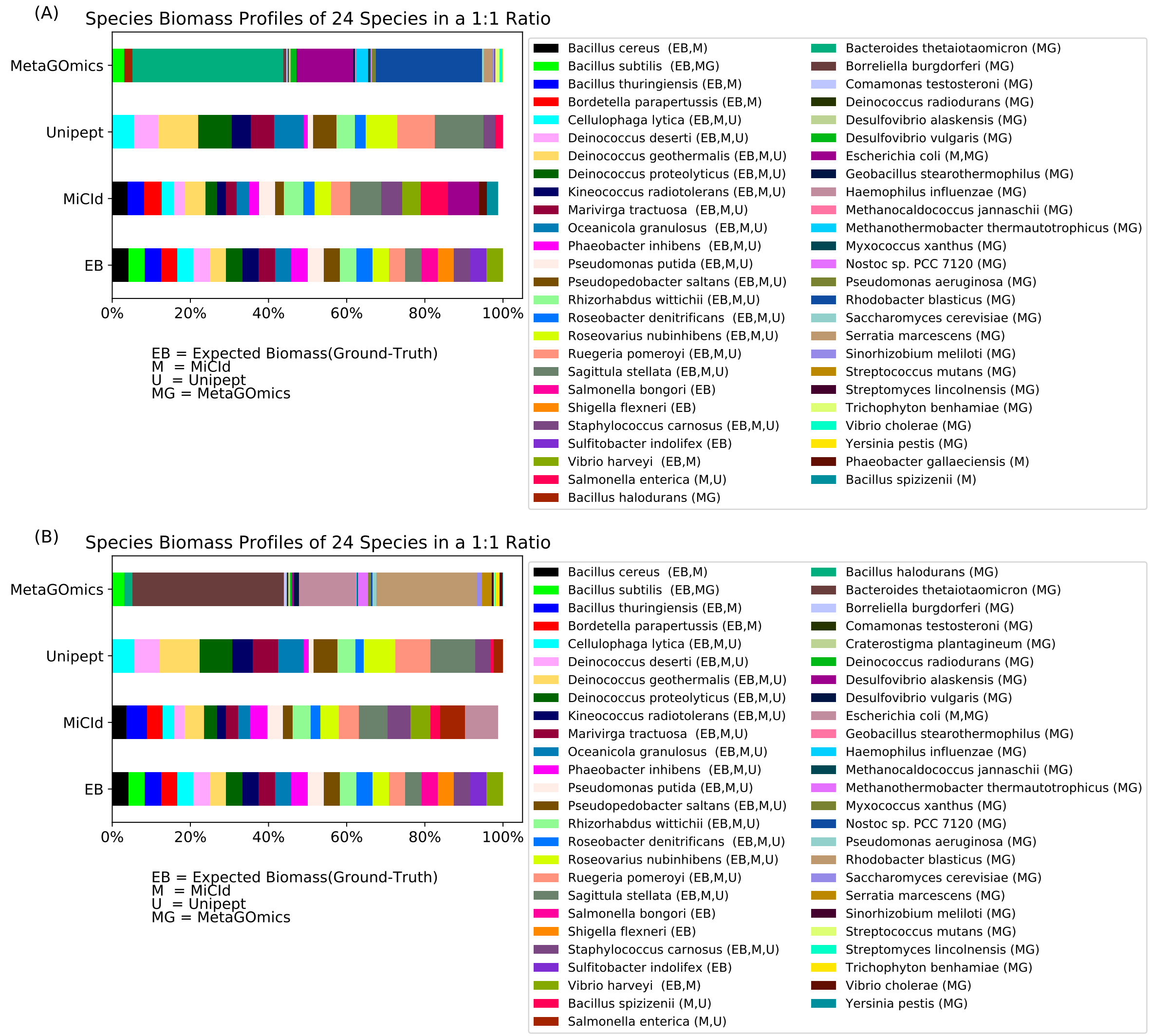


Figure S4: Comparison of species level biomass profiles reported MiCId and Unipept. Panels (A)-(B) presents stacked bar plots for a sample consisting of 24 species: Bacillus cereus, Bacillus subtilis, Bacillus thuringiensis, Bordetella parapertussis, Cellulophaga lytica, Deinococcus deserti, Deinococcus geothermalis, Deinococcus proteolyticus, Kineococcus radiotolerans, Marivirga tractuosa, Oceanicola granulosus, Phaeobacter inhibens, Pseudomonas putida, Pseudopedobacter saltans, Rhizorhabdus wittichii, Roseobacter denitrificans, Roseovarius nubinhibens, Ruegeria pomeroyi, Sagittula stellata, Salmonella bongori, Shigella flexneri, Staphylococcus carnosus, Sulfitobacter indolifex, and Vibrio harveyi in a 1:1 ratio (data file 20-21).

### Figure S5

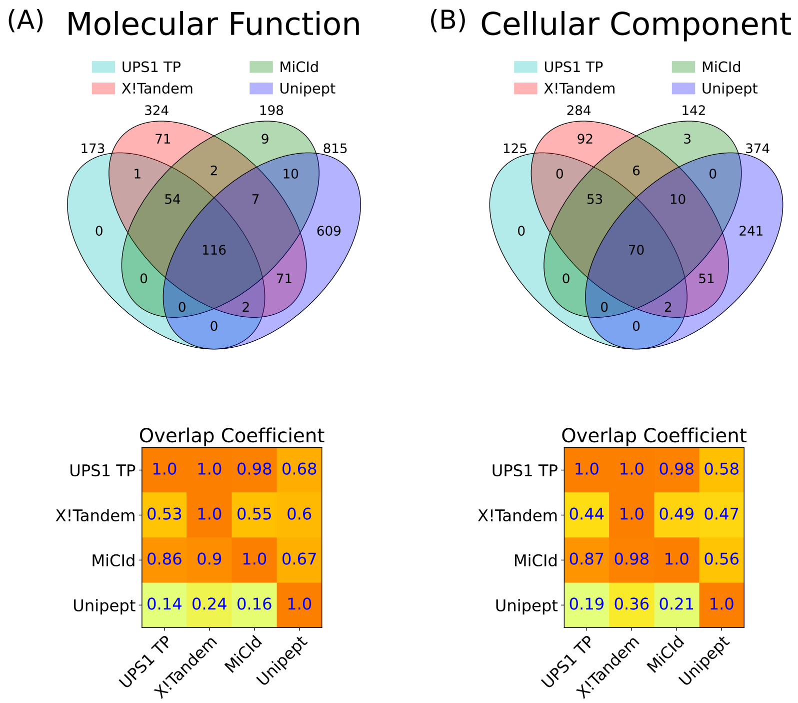


Figure S5: Assessment of GO term identification sensitivity and the proportion of false discoveries. Panels (A)-(B) present Venn diagrams and an overlap coefficient matrix for the GO terms reported by X!Tandem, MiCId, and Unipept. Panel (A) also includes the GO terms from the gold standard for the 48 human proteins in the UPS1 protein set (UPS1 TP). The Venn diagrams illustrate the number of GO terms identified by each method, as well as the number of terms co-identified by the methods. The overlap coefficient matrix values are calculated as the fraction of GO terms in the intersection between methods (corresponding to the row and column), divided by the total number of GO terms reported by the method in the row. When a GO term gold standard is used, as in panel (A)-(B) for UPS1 TP, the values in the first row of the matrix represent the sensitivity of the methods listed, while one minus the values in the first column indicates the proportion of false discoveries for those methods. The GO terms identified from the analysis of data file 7 are used in panels (A), and (B).

### Figure S6

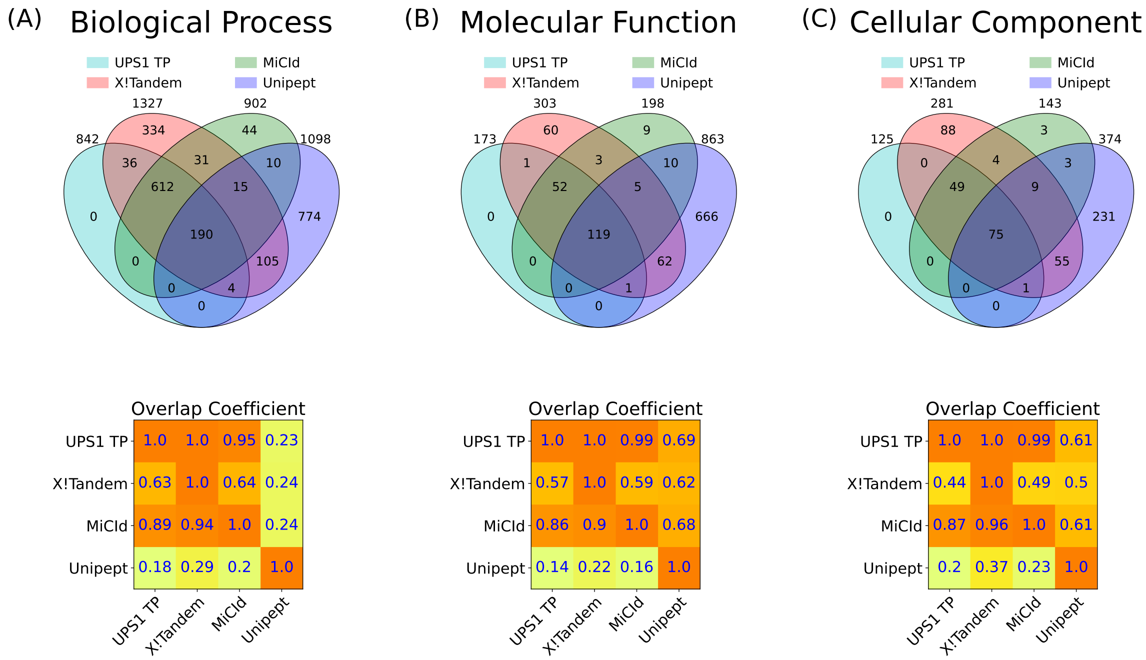


Figure S6: Assessment of GO term identification sensitivity and the proportion of false discoveries. Panels (A)-(C) present Venn diagrams and an overlap coefficient matrix for the GO terms reported by X!Tandem, MiCId, and Unipept. Panel (A) also includes the GO terms from the gold standard for the 48 human proteins in the UPS1 protein set (UPS1 TP). The Venn diagrams illustrate the number of GO terms identified by each method, as well as the number of terms co-identified by the methods. The overlap coefficient matrix values are calculated as the fraction of GO terms in the intersection between methods (corresponding to the row and column), divided by the total number of GO terms reported by the method in the row. When a GO term gold standard is used, as in panel (A)-(C) for UPS1 TP, the values in the first row of the matrix represent the sensitivity of the methods listed, while one minus the values in the first column indicates the proportion of false discoveries for those methods. The GO terms identified from the analysis of data file 8 are used in panels (A), (B), and (C).

### Figure S7

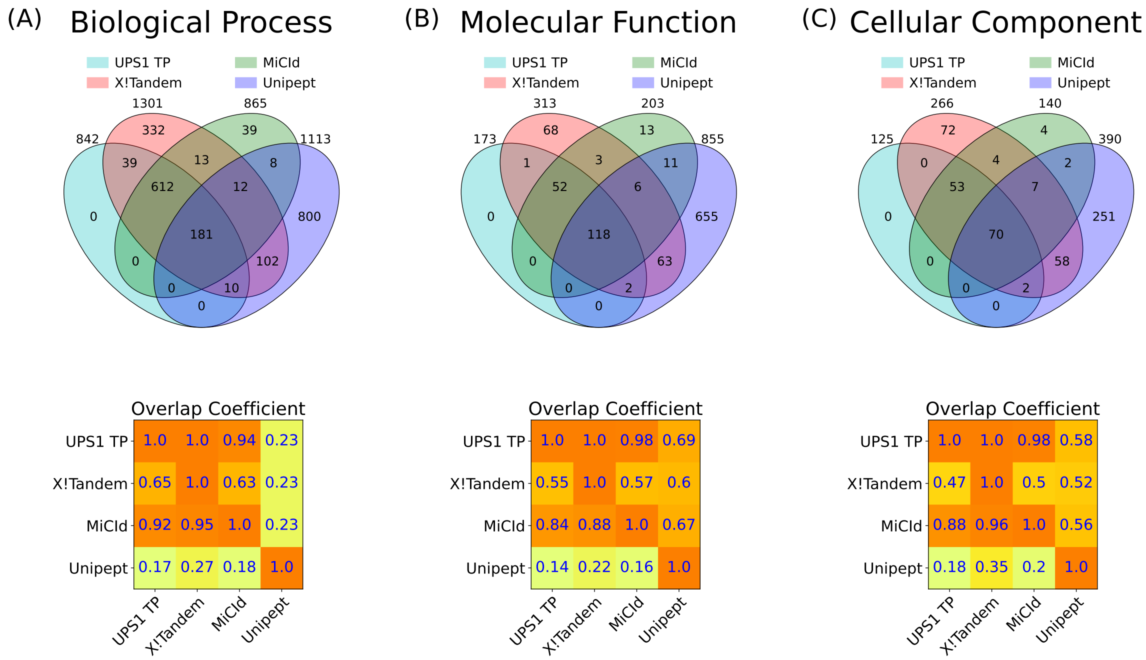


Figure S7: Assessment of GO term identification sensitivity and the proportion of false discoveries. Panels (A)-(C) present Venn diagrams and an overlap coefficient matrix for the GO terms reported by X!Tandem, MiCId, and Unipept. Panel (A) also includes the GO terms from the gold standard for the 48 human proteins in the UPS1 protein set (UPS1 TP). The Venn diagrams illustrate the number of GO terms identified by each method, as well as the number of terms co-identified by the methods. The overlap coefficient matrix values are calculated as the fraction of GO terms in the intersection between methods (corresponding to the row and column), divided by the total number of GO terms reported by the method in the row. When a GO term gold standard is used, as in panel (A)-(C) for UPS1 TP, the values in the first row of the matrix represent the sensitivity of the methods listed, while one minus the values in the first column indicates the proportion of false discoveries for those methods. The GO terms identified from the analysis of data file 9 are used in panels (A), (B), and (C).

### Figure S8

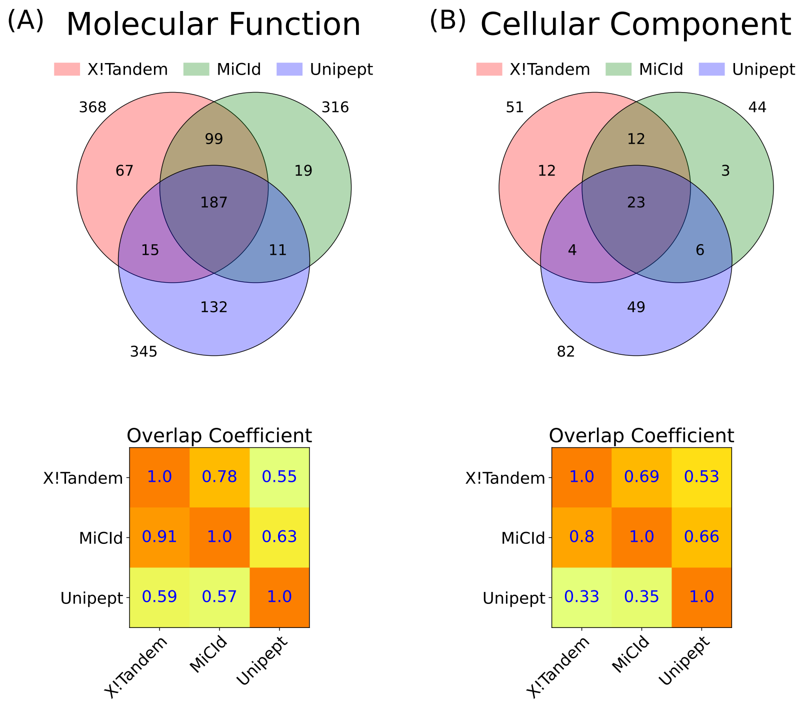


Figure S8: Assessment of GO term identification sensitivity and the proportion of false discoveries. Panels (A)-(B) present Venn diagrams and an overlap coefficient matrix for the GO terms reported by X!Tandem, MiCId, and Unipept. In panels (A)-(B), where a GO term quasi-gold standard is used (based on the GO terms reported by X!Tandem), the magnitude of the values in the first column reflects the performance of the methods listed in that column. The GO terms identified from the analysis of data file 10 are used in panels (A), and (B).

### Figure S9

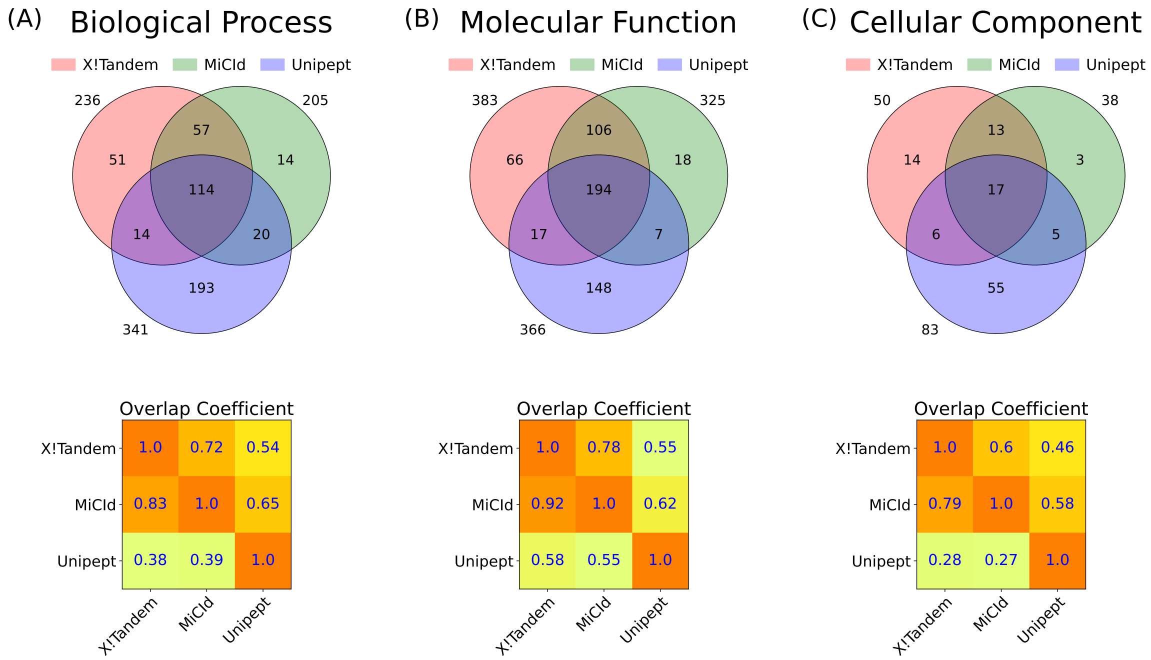


Figure S9: Assessment of GO term identification sensitivity and the proportion of false discoveries. Panels (A)-(C) present Venn diagrams and an overlap coefficient matrix for the GO terms reported by X!Tandem, MiCId, and Unipept. In panels (A)-(C), where a GO term quasi-gold standard is used (based on the GO terms reported by X!Tandem), the magnitude of the values in the first column reflects the performance of the methods listed in that column. The GO terms identified from the analysis of data file 11 are used in panels (A), (B), and (C).

### Figure S10

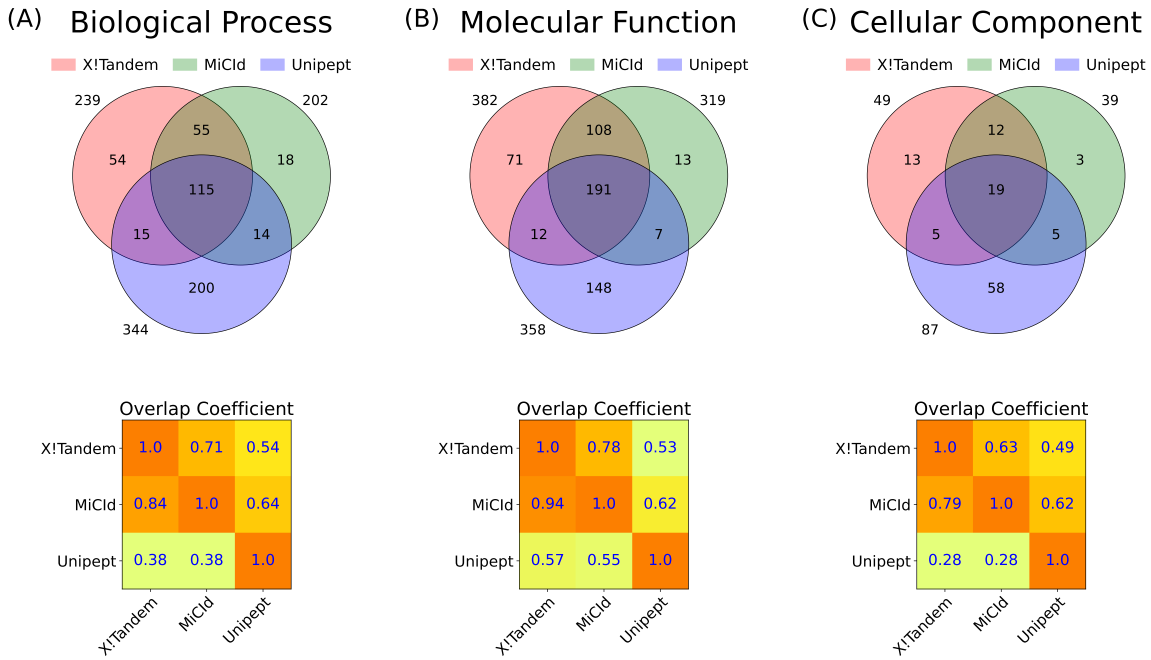


Figure S10: Assessment of GO term identification sensitivity and the proportion of false discoveries. Panels (A)-(C) present Venn diagrams and an overlap coefficient matrix for the GO terms reported by X!Tandem, MiCId, and Unipept. In panels (A)-(C), where a GO term quasi-gold standard is used (based on the GO terms reported by X!Tandem), the magnitude of the values in the first column reflects the performance of the methods listed in that column. The GO terms identified from the analysis of data file 12 are used in panels (A), (B), and (C).

### Figure S11

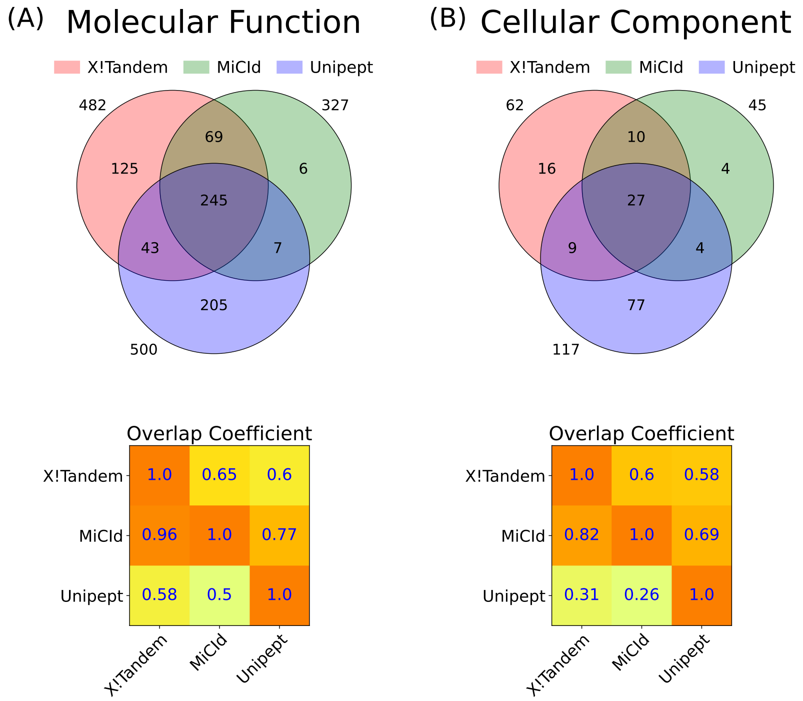


Figure S11: Assessment of GO term identification sensitivity and the proportion of false discoveries. Panels (A)-(B) present Venn diagrams and an overlap coefficient matrix for the GO terms reported by X!Tandem, MiCId, and Unipept. In panels (A)-(B), where a GO term quasi-gold standard is used (based on the GO terms reported by X!Tandem), the magnitude of the values in the first column reflects the performance of the methods listed in that column. The GO terms identified from the analysis of data file 19 are used in panels (A), and (B).

### Figure S12

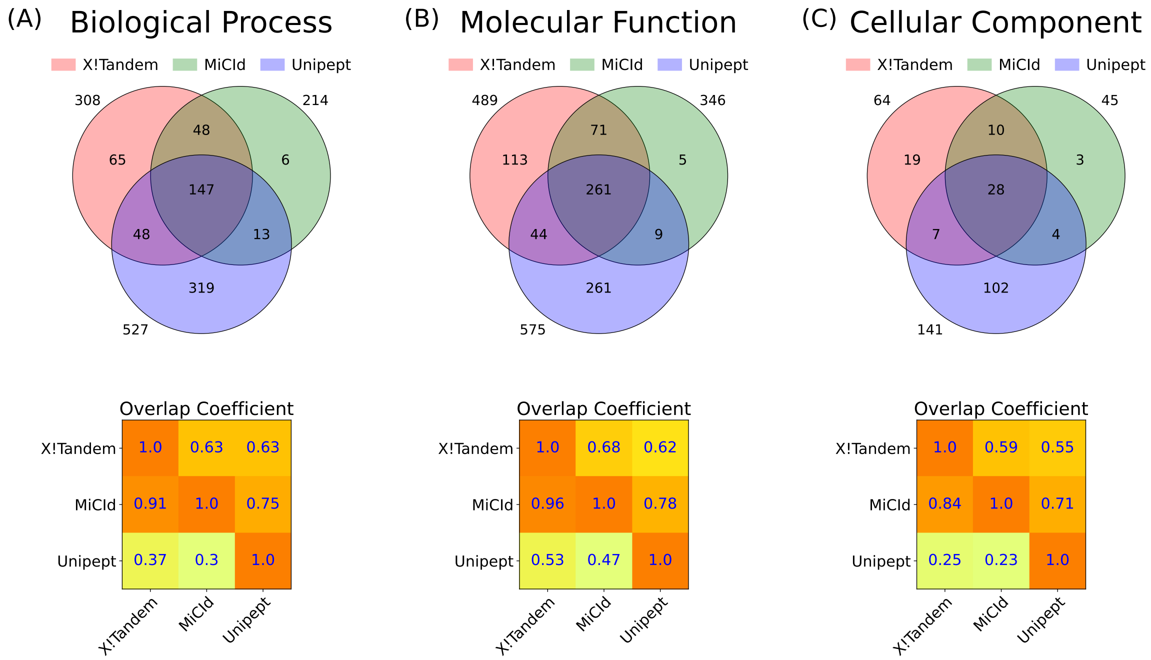


Figure S12: Assessment of GO term identification sensitivity and the proportion of false discoveries. Panels (A)-(C) present Venn diagrams and an overlap coefficient matrix for the GO terms reported by X!Tandem, MiCId, and Unipept. In panels (A)-(C), where a GO term quasi-gold standard is used (based on the GO terms reported by X!Tandem), the magnitude of the values in the first column reflects the performance of the methods listed in that column. The GO terms identified from the analysis of data file 20 are used in panels (A), (B), and (C).

### Figure S13

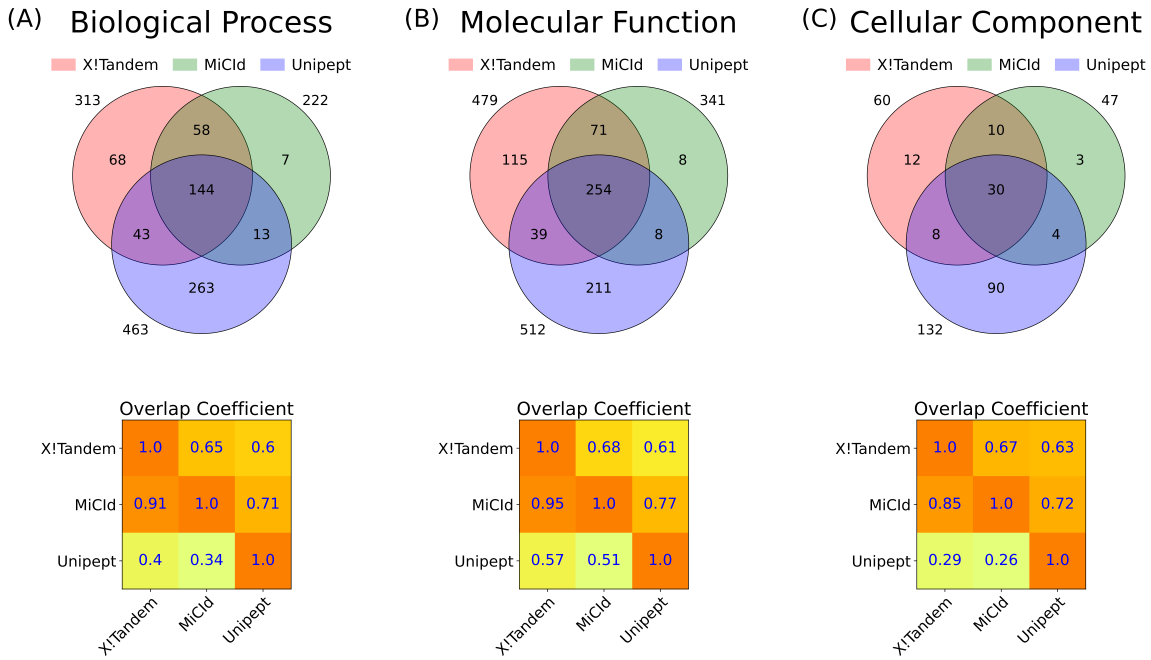


Figure S13: Assessment of GO term identification sensitivity and the proportion of false discoveries. Panels (A)-(C) present Venn diagrams and an overlap coefficient matrix for the GO terms reported by X!Tandem, MiCId, and Unipept. In panels (A)-(C), where a GO term quasi-gold standard is used (based on the GO terms reported by X!Tandem), the magnitude of the values in the first column reflects the performance of the methods listed in that column. The GO terms identified from the analysis of data file 21 are used in panels (A), (B), and (C).

### Figure S14

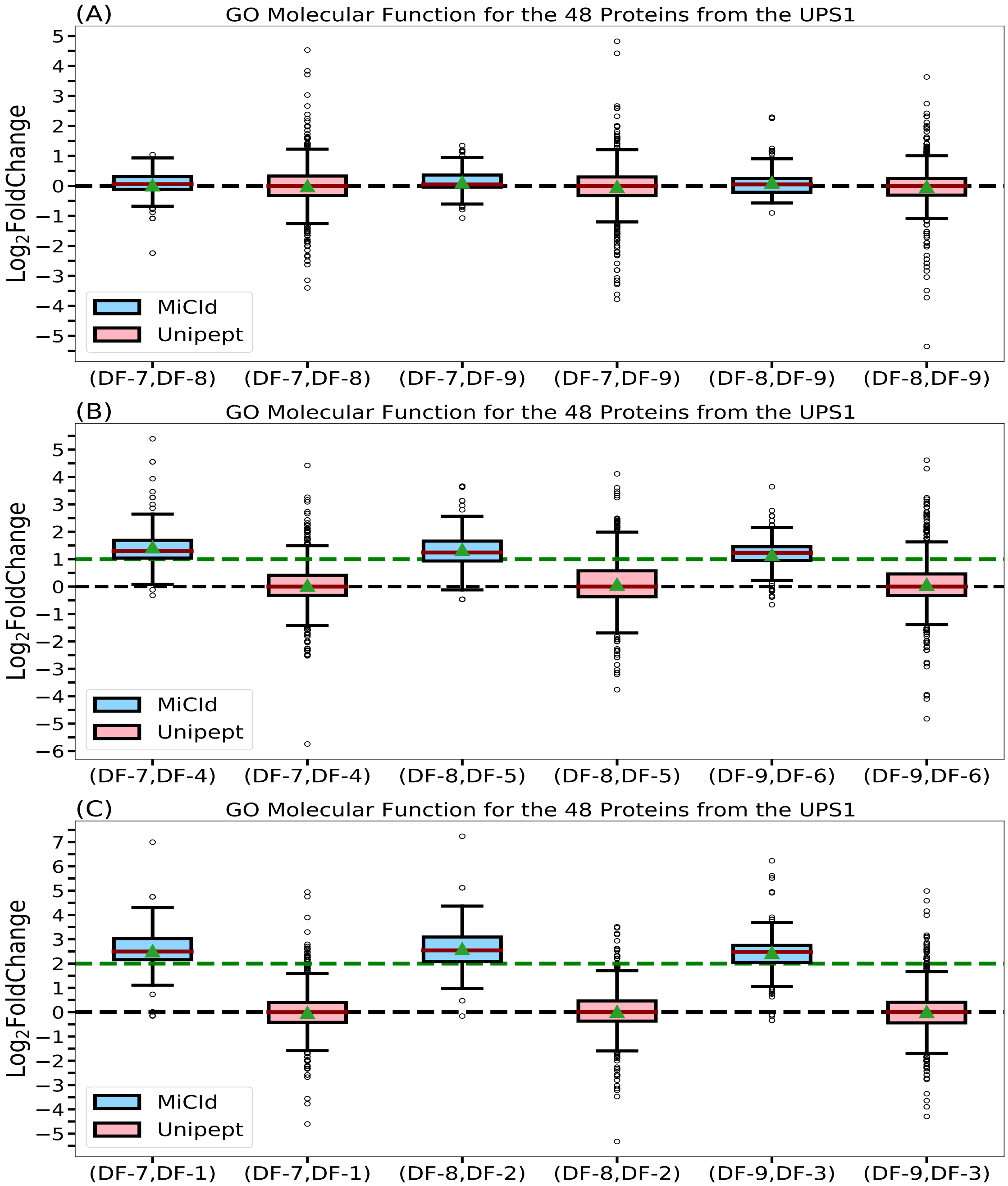


Figure S14: Assessment of GO term abundances computed by MiCId and Unipept. Panels (A), (B), and (C) present box-whisker plots for the $\log_2$ fold change between samples with varying amounts of the 48 human proteins from the UPS1 protein set, at 1:1, 2:1, and 4:1 ratios, respectively. In each panel, an accurate and precise biomass estimation method should result in a box-whisker plot centered around the expected values of 0, 1, and 2, respectively, with a narrow width, indicating minimal deviation from the expected biomass. The $\log_2$ fold change in GO term abundances is derived from the analysis between the data files (DF-i, DF-j), as indicated by the x-axis labels for each box-whisker plot.

### Figure S15

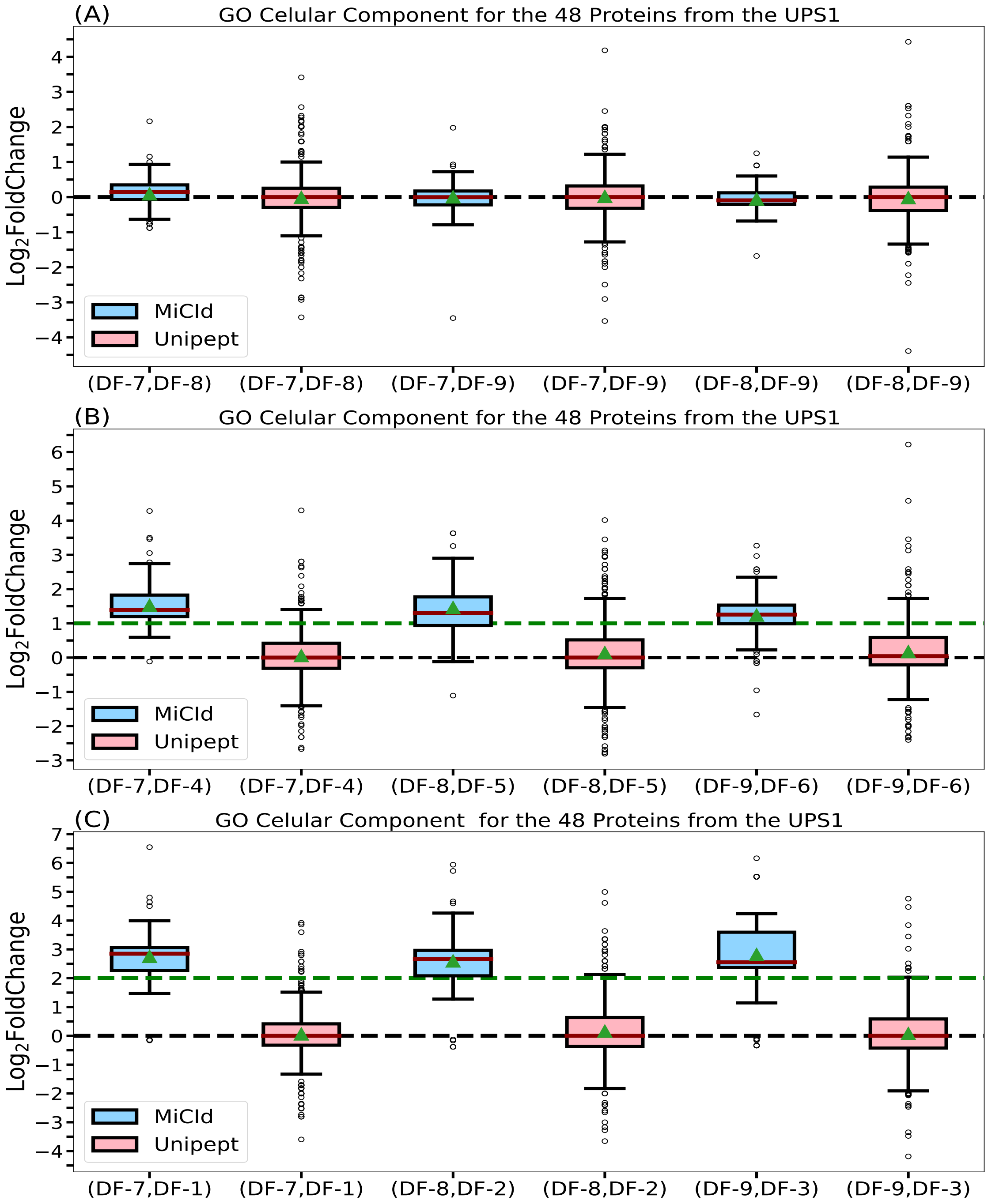


Figure S15: Assessment of GO term abundances computed by MiCId and Unipept. Panels (A), (B), and (C) present box-whisker plots for the $\log_2$ fold change between samples with varying amounts of the 48 human proteins from the UPS1 protein set, at 1:1, 2:1, and 4:1 ratios, respectively. In each panel, an accurate and precise biomass estimation method should result in a box-whisker plot centered around the expected values of 0, 1, and 2, respectively, with a narrow width, indicating minimal deviation from the expected biomass. The $\log_2$ fold change in GO term abundances is derived from the analysis between the data files (DF-i, DF-j), as indicated by the x-axis labels for each box-whisker plot.

### Figure S16

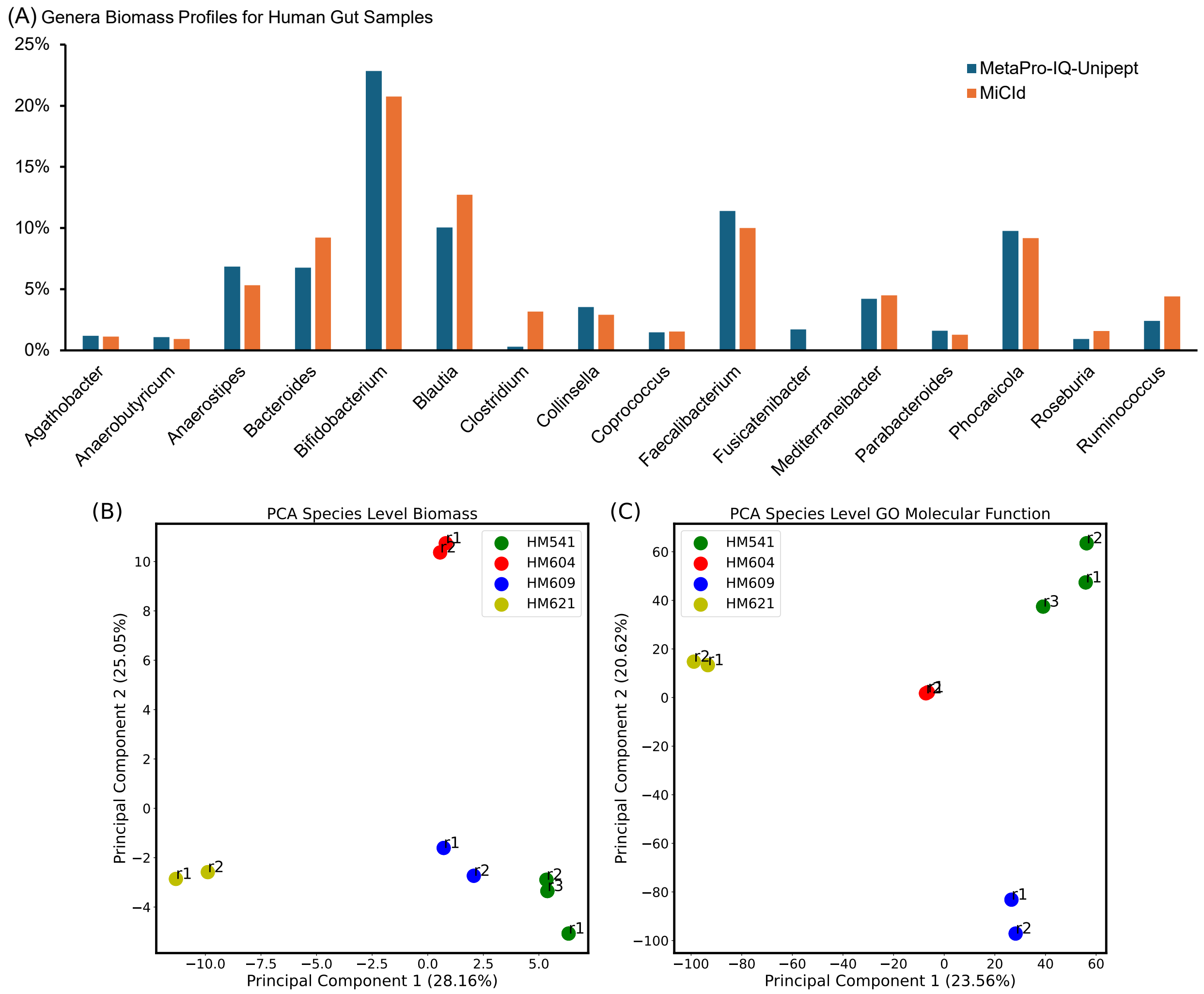


Figure S16: Data analysis results for the human gut microbiome dataset. Panel (A) presents a bar plot depicting the average genera biomass composition for human gut samples, derived from the results of MiCId and MetaPro-IQ-Unipept. Only genera identified by at least one of the methods with an average biomass of 1% or greater are included. The plot shows a high degree of similarity between the two methods, with a correlation coefficient of 0.96 between the average genera biomasses obtained from MiCId and MetaPro-IQ-Unipept. Panels (B) and (C) display PCA plots for species-level biomass abundances and GO term molecular function abundances, respectively. All the confidently identified species ($E$-value $\leq$ 0.01) and GO terms are included in the PCA plots. The abundances used in the PCA plots were computed using the proposed EM algorithm implemented in MiCId workflow, logit-transformed with a base-2 logarithm, and standardized by subtracting the mean and dividing by the standard deviation. The PCA plots demonstrate the high reproducibility of the EM algorithm in estimating biomass abundances for taxa and biological functions, with technical replicates of the same sample clustering significantly closer together than replicates from different samples. The results for technical replicates $r_1$ and $r_2$ of sample HM604 are difficult to distinguish in panel (B) because the data points are nearly 100% superimposed. In the PCA plots, technical replicates are represented by circular points in the color of the corresponding sample and labeled with $r_i$.

### Figure S17

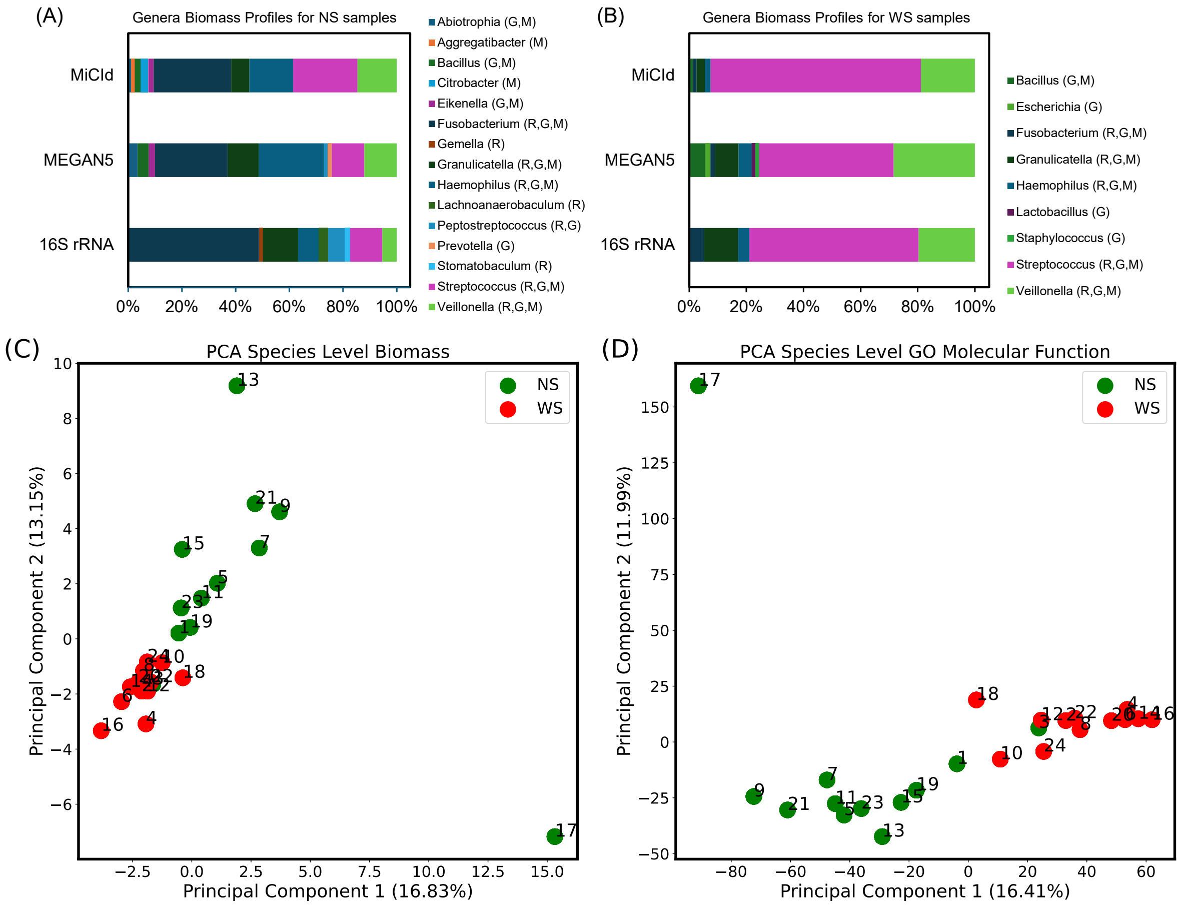


Figure S17: Data analysis results for the human oral microbiome dataset, consisting of 12 samples incubated in biofilm reactors under two conditions: with sugar (WS) and without sugar (NS). Panels (A) and (B) show stacked bar plots displaying the average genera biomass composition for human oral microbiome samples, based on MiCId, MEGAN5, and 16S rRNA results. Only genera with an average biomass of 1% or greater are included. These plots highlight a distinct difference in genera composition between the NS and WS samples and demonstrate notable consistency in the average biomass profiles calculated for MiCId, MEGAN5, and 16S rRNA within both conditions. Panels (C) and (D) depict PCA plots for species-level biomass abundances and GO term molecular function abundances, respectively, both computed using the proposed EM algorithm implemented in the MiCId workflow, logit-transformed with a base-2 logarithm, and standardized by subtracting the mean and dividing by the standard deviation. The PCA plots reveal clear separation of NS and WS samples along the first principal component, except for samples 730NS and 733NS, marked by green circles and labeled as 1 and 3. In the PCA plots, for clarity, NS samples are represented by green circles (odd-numbered labels 1, 3,· · · , 23), while WS samples are represented by red circles (even-numbered labels 2, 4,· · · , 24).
